# Skin-derived G-CSF activates pathological granulopoiesis upon psoriasis

**DOI:** 10.1101/2025.10.10.681054

**Authors:** Tomson Kosasih, Tatsuya Morishima, Sohyeon Lee, Jungyeon Yoon, Kanako Wakahashi, Pilhan Kim, Aiko Sada, Hitoshi Takizawa

## Abstract

Psoriasis is an inflammatory skin disease initiated by environmental triggers and driven by disruption of T cell cytokine network in the cutaneous milieu. The fact that complete resolution of disease by targeting key inflammatory cytokines remains challenging indicates a contribution of other immune cells to the pathogenesis. Here, we study the role of neutrophils in psoriasis, the first-line innate immune defender that is short-lived but mobile and infiltrate into various tissues. We found that upon psoriasis induction, skin-resident endothelial cells are activated to produce G-CSF which activates emergency granulopoiesis in bone marrow and induces cutaneous infiltration and accumulation of neutrophil that are functionally overactive. Depletion of neutrophils or blockage of psoriasis-driven granulopoiesis by respective neutralizing antibodies results in reducing cutaneous neutrophil burden and mitigating psoriasis pathogenesis. This mechanism might be conserved in human psoriasis as confirmed by public RNA-seq database. Our findings uncovered and detailed the pathological crosstalk between skin and BM in psoriatic inflammation, proposing a potential therapeutic approach targeting cross-organ communication.

## INTRODUCTION

Psoriasis is an inflammatory skin disease initiated by environmental triggers such as infections, air pollutants, and excessive sunlight exposure, progressively abolishing tissue integrity and aggravate the pathologies overtime.^1^ At its core, psoriasis is an immunological disorder driven by disruption of cytokine network, particularly IL-17 overtone caused by hyperactive T-helper 17 (Th17) in the cutaneous milieu. Although Th17-based immunopharmacological rewiring mitigates the disease, complete resolution remains largely unachievable,^2–5^ indicating the likely contribution of other immune cell subsets.

A hallmark of psoriasis is the formation of microabscesses caused by neutrophil infiltration.^6^ Neutrophils are the first-line innate immune defender which respond quickly and effectively to broad spectrum of antigens.^7^ Upon systemic infection, granulopoiesis is rapidly activated in the BM, shifting hematopoiesis toward neutrophil generation, often referred to as emergency granulopoiesis, in order to clear pathogen.^8^ However, little did it known whether local inflammation in peripheral organ can elicit hematopoietic response in distal bone marrow, and if so, what is the underlying mechanism to mediate cross-organ communications.

Tissue and circulating neutrophils are naturally short-lived and constantly replenished through *de novo* granulopoiesis in the bone marrow (BM). IL-1, IL-3, IL-6, granulocyte-macrophage colony-stimulating factor (GM-CSF), and granulocyte-colony stimulating factor (G-CSF) are key cytokines that differentiate hematopoietic stem progenitor cells (HSPCs) into granulocyte lineage and determines cell pool in a systemic manner. Especially G-CSF is shown to be secreted by BM-resident endothelial cells upon its Toll-like receptors (TLRs) sensing and drive emergency granulopoiesis.^9–11^ Given the excess accumulation of short-living neutrophil in the skin upon psoriasis, we hypothesized that skin would initiate long-distance communication to the BM possibly through a blood-circulating signal, and activate emergency granulopoiesis program to meet the local demand in inflamed skin.

Here, using an inducible psoriasis mice model,^12^ we demonstrated that psoriatic skin insult induced cutaneous neutrophil infiltration and G-CSF production exclusively by the skin endothelial cells (ECs) which activates bone marrow emergency granulopoiesis. Transcriptomic analysis reveals that skin-infiltrating neutrophils were functionally overactive. Psoriasis-driven emergency granulopoiesis was lessen by anti-GCSF neutralization resulting in the lowered cutaneous neutrophil burden leading to disease mitigation. This mechanism reflects human psoriasis setting as RNA-seq public data reanalysis reveals the analogous phenomenon. Our findings uncovered and detailed the pathological crosstalk of skin-BM axis in psoriatic inflammation.

## METERIALS AND METHODS

### Mice

C57BL/6 (CD45.2) founder mice were purchased from Japan SLC, Inc. and bred/maintained at the Center for Animal Resources and Development at Kumamoto University. Ly6G^Cre^; Rosa26^tdTom^ (CD45.2)^13^ mice were purchased from The Jackson Laboratory. All the animal experiments were performed in female mice age 8–12-week-old. Animal experiments were conducted under the approval of Animal Care and Use Committee of Kumamoto University.

### Animal Treatment

Psoriasis was induced by daily topical application of 62.5mg Imiquimod (IMQ) cream (Beselna, Mochida Pharmaceutical) to the pre-shaved and hair-removed 2.5x2.5cm dorsal skin for four consecutive days. Psoriasis clinical score (erythema, scaling) was measured according to the previous studies.^12,14^ In short, the skin was observed and clinically scored in a scale representing none (0), mild (1), moderate (2), significant (3), and severe (4). Histological analysis of dorsal skin was performed with Hematoxylin/Eosin staining and images were taken with light microscope, followed by analysis in Image J 1.53t.

Anakinra (KINERET, Biovitrium) (37μg) was intraperitoneally (i.p.) administered daily starting at 1 day before psoriasis induction (-1d) till 1 day before mice analysis (3d). Neutrophil was depleted by daily i.p. administration of 50μg anti-Ly6G (1A8, BioXCell) between -2d and 3d. The same dose of IgG-isotype (2A3, BioXCell) was given to control mice following the same regimen. Anti-G-CSF (MAB414, R&D Systems) or IgG-isotype control (AF007, R&D Systems) (10-20μg) was i.p. injected daily from 0d to 3d 30 minutes prior to IMQ topical association.

### Cytokines measurement

The peripheral blood was harvested retro-orbitally using untreated glass capillary (Hirschmann) and sat at RT for 30 min until clotted. Subsequently, the serum was optimally separated by centrifuging (2000xg; 10min; 4°C), collected, and snap frozen in liquid nitrogen before storage at -80°C till analysis. G-CSF and other cytokines in the serum were measured using Mouse G-CSF ELISA kit (Proteintech) and flowcytometry-based multi-cytokine measurement Legendplex^TM^ Mouse Inflammation Panel (Biolegend), respectively, according to the manufacture’s instruction.

### Flowcytometry analysis

To prevent contamination of circulating cells into tissue, intracardiac PBS perfusion was conducted before tissue collection. Immediately after the procedure, 4cm^2^ skin was harvested and subsequently cleared from the remnant hair and dead skin (scale) gently with scalpel. The resultant skin was minced in a digestion buffer containing 62.5µg/mL Liberase^TM^ (Roche), 50µg/mL DNAse (Roche), 10mM HEPES (Nacalai Tesque) in RPMI-1640 medium (Wako), followed by incubation with gentle rocking at 37°C for 2h. An equal volume of pre-warmed 0.1% BSA/HBSS (Merck/Sigma) was added to the mixture and mechanical tissue homogenization was performed using Gentle MACS^TM^ (program C_01) to prepare a homogeneous single cell suspension. Subsequently, the skin homogenate was filtered stepwise with 100μm, 70µm, and 40µm strainer (BD Biosciences), and centrifuged at 400 x g for 5 min 4°C to harvest skin cells in a pellet and instantly resuspend in 2% FBS 0.2mM EDTA in PBS (FACS buffer). To reduce non-specific immunostaining, Fcγ blocking was performed by incubating the cell suspension with CD16/32 (93) for 15min in RT prior to antibody-staining with fluorophore-conjugated antibodies (30min on ice) against the following markers: CD45.2 (104), CD19 (6D5), CD11b (M1/70), Ly6C (HK1.4), Ly6G (1A8), CD31 (390), Ter119 (TER-119), CD45 (30-F11), CD49f (GoH3), CD207 (4C7), CD101 (Moushi101), CD11c (N418), Sca-1 (D7), and CD34 (RAM34).

For BM cells, cells were collected by grinding two hindleg bones (tibiae and femurs) using a mortar and pestle in FACS buffer before treatment with ammonium-chloride-potassium buffer (0.15mM NH_4_Cl, 1.0mM KHCO_3_, 0.1mM Na_2_EDTA) to remove red blood cells. The BM single cell suspension was preincubated with CD16/32 (93) in RT for 15min, prior to staining with the fluorophore conjugated antibodies/reagents against CD45 (30-F11), CD11b (M1/70), Ly6G (1A8), IL7ra (A7R34), c-Kit (2B8), Sca-1 (D7), CD16/32 (93), CD34 (RAM34), CD48 (HM48-1), CD150 (TC15-12F.2), CD135 (A2F10), and streptavidin-conjugated antibody for 30-60 min on ice. For HSPC analysis, additional staining with biotinylated-antibodies against Ter119 (TER-119), CD3 (145-2C11), CD4 (GK1.5), CD8 (53-6.7), NK1.1 (PK136), B220 (RA3-6B2), CD11b (M1/70), Gr-1 (RB6-8C5) was performed before fluorophore-antibodies association, while Fcr blocking with CD16/32 (93) was omitted.

For all FACS analysis, dead cells were stained with Hoechst33342 or Propidium Iodide shortly before analysis, and analysis was done within 4h after. Samples were analyzed on FACS Symphony A3 or FACS Aria IIIu (BD Biosciences). All the data were further analyzed using Flowjo10.10.0.

### Quantitative RT-PCR

For tissues, 50 mg of the snap frozen sample was pulverized using an ultra-cold conditioned (liquid nitrogen soaked) metal mortar and pestle (Tokken, inc.). RNA was isolated from the tissue homogenate with RNeasy Mini Kit (Qiagen) and reverse transcribed into cDNAs with reverse transcriptase containing reagents mix (Takara). The resultant cDNA was subjected to RT-PCR with the respective primers (Supplemental information), and the relative expression of gene of interest was quantified relative to housekeeping genes (*Gapdh or B2m*).

For cells, 100 cells were FACS-sorted into the designated 96 well-plate containing cell lysis buffer and subjected to cDNA synthesis according to the detailed protocol described in the previous study adapted for 100 cells processing, followed by RT-PCR.^15^

### Skin Endothelial Cell Culture and Stimulation

Dorsal skin from healthy mice were harvested and digested into single cell suspension in prior to fluorophore conjugated antibody staining for immunophenotypic endothelial cell (CD45^-^ Ter119^-^ CD31^+^ Sca-1^+^). Sixty thousand endothelial cells were sorted into a flat bottom 96-well plate pre-coated with 0.2% gelatin. The sorted cells were recovered in 50% FBS, 100 unit/mL Penicillin G, 100 µg/mL Streptomycin (Wako), 2mM Glutamax (Gibco), and 1x NEAA (Gibco) in DMEM (all Thermo Fisher Scientific) at 37°C, 5% O_2_ for 36h. Subsequently, the media is substituted with stimulating media containing 20% FBS, 100unit/mL Penicillin G, 100µg/mL Streptomycin (Wako), 2mM Glutamax (Gibco), and 1x NEAA (Gibco) with 5% DMSO (Wako) as a control or 50 µg/mL IMQ (Sigma) in DMEM at 37°C, 5% O_2_ for another 48h. Finally, media was harvested for G-CSF measurement by ELISA (Proteintech).

### Three dimensional (3D) intra-vital imaging

An image acquisition window to image the same skin site was installed in the mouse’s dorsal site and mice were i.v. injected with fluorophore-conjugated CD31 antibody. 3D images were acquired daily from d0 to d2.

From the raw 3D-images, the blood vessels (CD31^+^) were transformed into Surface, and neutrophils (Ly6G^+^) were transformed into spots. The distribution of spot at 0-10, 10-20, 20-30, 30-40, 40-50, and >50µm distance to the surface was quantified. At the same time, random spots were generated using Python based module tifffile ver. 2025.3.30 to create 3D tiff files containing randomly distributed spheric voxels with x/y/z of 2.5µm to be overlaid into the original 3D imaging file. The amount of random spot per stacked files is equal to the number of phenotypic neutrophils per 3D image, transformed into spot, and follow the same distance measurement to surface as was done with neutrophil spots. Image processing and analysis was done in Imaris 8 (Oxford Instruments).

### RNA sequencing

One thousand neutrophils (CD45^+^CD11b^+^Ly6C^mid^Ly6G^+^) were FACS-sorted from dorsal skin and processed to RAMDA-seq.^16^ In short, the cells are disintegrated using lysis-buffer (Roche) and the whole RNAs were reversed transcribed into first strand cDNAs (Takara Bio Inc.). Using Klenow Fragments (3’-5’ exo; New England Biolabs), the double strand cDNAs was created which were magnetically purified with AMPure XP beads (Beckman Coulter) and further processed using Nextera XT DNA sample Prep kit to generate a library (Illumina). The quality and quantity of the library was measured with Agilent 4150 TapeStation (Agilent Technologies, Santa Clara, CA) and GenNext® NGS library quantification kit (Toyobo). Upon confirmation of adequate quality and quantity of the library, it was sequenced using NovaSeq X system (Illumina). Quality check and trimming of single-end sequences were completed with the trim_galore (version 0.4.3) package. Quality and length parameters were quality 30 and length 30. Filtered sequences were processed to remove mitochondrial mRNA with the SortMeRNA (version 2.1.1) and were aligned to mouse reference sequences (GRCm39) from (http://www.genecodesgenes.org/mouse/release.html) with ultrafast RNAseq aligner STAR (version 2.7.10b; cite Dobin et al, 2012). All the aligned bam files were used with mouse GFF annotation files (GRCm39) as input into the featureCounts program from the Subreads program (version 2.1.1) to count the raw reads for each gene and sample, and to create a gene count matrixDES. To calculate differentially expressed genes, the DESeq2 (version 1.30.1) package was used in R. The gene ontology (GO) analysis was performed in GSEA 4.4.0 software following default set parameters with customization including m5.all.v2025.Mm.symbols.gmt (https://www.gsea-msigdb.org/gsea/msigdb/mouse/genesets.jsp?collection=M5), Gene sets database reference, 1000 Number of permutations, gene_sets Permutation type, and Signal2Noise metric for ranking genes.^17,18^ Gene expression levels were quantified using Z-scores, with positive values indicating upregulation and negative values indicating downregulation.

### Re-analysis of human psoriasis RNA-seq data

The gene-expression dataset GSE54456^19^ is a skin bulk-RNA-seq data which has been normalized by reads per kilobase of exon per million mapped reads (RPKM) publicly available in Gene Expression Omnibus (GEO) repository. The data contains RNA expression level from 82 normal and 92 psoriatic human skin biopsies. The raw data from GEO was imported and incorporated to its metadata (sample group, methods, etc.) using GEOquery v.2.76.0 package. Subsequently, UMAP clustering was done using umap v.0.2.10.0 and plotly 4.11.0 packages. Those data processing was done in R-studio build 394. The expression value of the gene of interest was extracted to create customized bar charts in Prism version 10.

The GSE173706 single-cell RNA-seq dataset was obtained from the GEO public repository. ^20^ Each sample was filtered to remove cells with >5% mitochondrial content and potential doublets (2% of total cells). Filtered samples were then integrated, and expression counts were log-transformed to 10^4^ per cell. Clustering analysis was done in scvi-tools v.1.2.0 and was used to identify cellular clusters which were annotated based on the original study.^20^ Gene expressional level mapping and cell type frequencies were calculated using scanpy v.1.10.3.

### Statistical analysis

The Shapiro-Wilk test was performed to assess the data distribution, identifying it as normal (*p*≥0.05) or skewed (*p*<0.05). For dataset containing 2 groups, either an unpaired T-test (normal distribution) or a Mann-Whitney (skewed distribution) statistical test was performed. For dataset containing more than 2 groups, one-way ANOVA (normal distribution) or Kruskal-Wallis (skewed distribution) with Dunnet or Dunn test for its multiple group comparison was done. Prism version 10 was used to generate graphs and to perform statistical analysis in this study with the star count annotation reflecting significance level as: \**p*<0.05; \*\**p*<0.01; \*\*\**p*<0.001; \*\*\*\**p*<0.0001.

## RESULTS

### Psoriasis induces progressive infiltration of neutrophil to skin

We employed a well-established murine psoriasis model by topical application of a TLR7 agonist, imiquimod (IMQ) or vaseline control (Vas) for four consecutive days (Figure 1A). This treatment induced acute psoriasis-like manifestations, including scaling, erythema, and epidermal thickening at the application site (Figure 1B and S1A), accompanied by increased expression of dermatitis-associated cytokines such as *Il17a, Il17f, Il22, and Il6* (Figure S1B).^12,21^ Flow cytometric analysis revealed that, among skin myeloid subsets, neutrophils were notably expanded following IMQ treatment (Figure S1C). Their numbers increased progressively, reaching approximately 10-fold by day 4 (4d), whereas Ly6C^+^ and Ly6C^-^ myeloid cells are unchanged (Figure 1C). Consistently, the neutrophil-attracting chemokines *Cxcl1, Cxcl2, and Cxcl5* were upregulated by 10-100 fold in the psoriatic skin, while *Ccl2,* a major monocyte chemoattract, remained unchanged (Figure S1D).^22,23^ Neutrophil infiltration was restricted to IMQ-treated skin, as it was absent in untreated skins of the same animals (Figure S1E).

**Figure 1.**
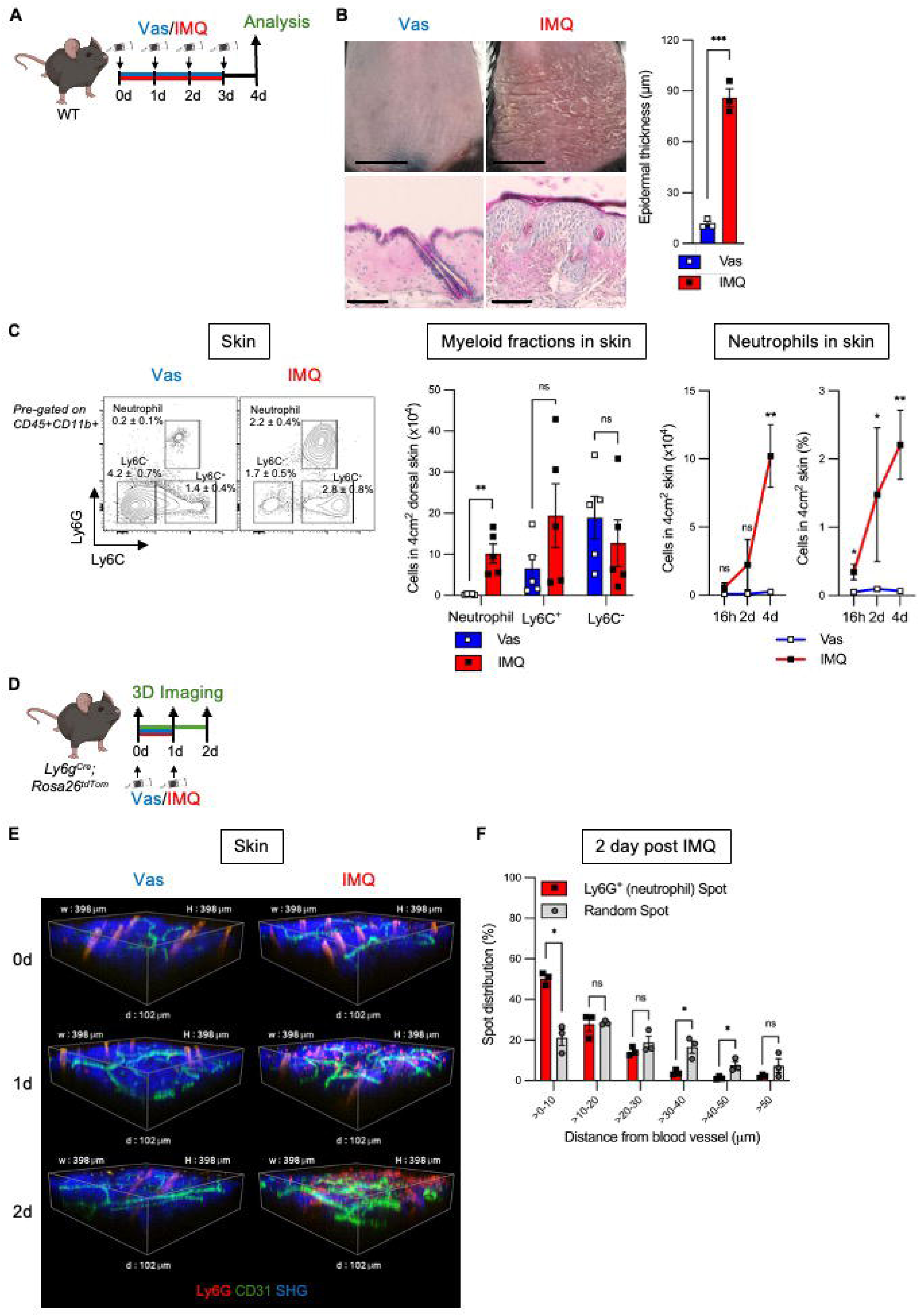
Psoriasis induces neutrophil migration and infiltration to the inflamed skin. **(A)** Experimental scheme of IMQ-induced psoriasis mouse model for results depicted in (B and C) (Vas, vaseline cream; IMQ, Imiquimod cream). **(B)** Clinical manifestation of psoriasis at 4d. Representative skin images (upper panel, shaved dorsal skin with 1cm scale bar; lower panel: HE histology with 100µm scale bar). Bar graph shows epidermal thickness (µm) in mice treated with Vas (n=3) or IMQ (n=3). **(C)** FACS analysis of myeloid cells in skin. Representative FACS plot pre-gated on CD45^+^CD11b+ cells. Number of myeloid cells (neutrophil, Ly6C^+^ and Ly6C^-^ cells) at 4d after treatment with Vas (n=5) or IMQ (n=5). Time-course kinetics of skin neutrophils at 16h, 2d, and 4d following topical application with Vas (n=3-5) or IMQ (n=3-5). **(D)** Experimental scheme of intravital microscopic skin imaging for results depicted in (E and F). **(E)** Representative time-course images of neutrophils (Ly6G^+^), blood vessel (CD31^+^), and collagen (SHG) at 0d, 1d and 2d post Vas/IMQ treatment. **(F)** Distribution of neutrophil localized from nearest blood vessel measure from 2d post IMQ compared to random spots (n=3). Data are pooled from ≥2 independent experiments. Each dot shown in the chart represents the measurement from 1 experimental subject and shown as mean±S.E. \**p*<0.05; \*\**p*<0.01; \*\*\**p*<0.001.

To further define neutrophil trafficking, we performed intravital 3D imaging in the dorsal skin of *Ly6G^Cre^;Rosa26^tdTom^* mice in which neutrophils are labeled with tdTomato.^13^ The analysis was done between 0d and 2d due to high autofluorescence of hair follicle in the psoriatic skin after 2d (Figure 1E and S1F). The CD31^+^ blood vessels expanded in parallel with neutrophil accumulation (Figure 1E and S1F). At baseline (0d), neutrophils were evenly distributed within 50µm distance from the nearest vessels. By 1d post psoriasis induction, ∼40% of them are localized within 0-10µm and another 40% within 10-20µm. About 50% of neutrophils were concentrated within 0-10µm and about 30% within 10-20µm by 2d, indicating progressive perivascular neutrophil clustering during psoriasis development (Figure 1F, S1G, and S1H).

Collectively, these results suggest that neutrophils migrate to the skin and are retained in close proximity to blood vessels during psoriasis.

### Skin neutrophils upregulate IL-17A *and* promote psoriasis progression

To study the functional role of neutrophil in psoriasis, we performed RNA-seq with neutrophils sorted from both control (Vas) and psoriatic (IMQ) skin at 4d post-induction. Principal component analysis revealed distinct transcriptional profiles that IMQ-treated neutrophils are differentiated from the closely-clustered Vas-treated neutrophils (Figure 2A). Neutrophils in inflamed sites are shown tissue damaging as they are metabolically active to facilitate intensification of reactive oxygen species and proteolytic enzymes production, besides their tendency to undergo Neutrophil Extracellular Trap (NET)osis.^24,25^ Gene ontology (GO) analysis revealed that neutrophils in the IMQ-treated skin didn’t have heightened metabolic activation compared to steady-state skin-resident neutrophils (Figure S2A), but rather show an increased sensing of neurotransmitter and sensory molecules. Instead, the phagocytic and the respiratory related activities were downregulated (Figure 2B).Since IL-17 is the key cytokine in psoriasis pathology^26^ and IL-17 secreting neutrophils are found in psoriatic lesion of patients,^27,28^ we reason that neutrophil might contribute to psoriasis inflammation by enhancing IL-17 cutaneous tone. Among all *Il17* subtypes checked in the RNA-seq data, IMQ-treated neutrophils specifically upregulate *Il17a* among all subtypes (Figure 2C). RT-PCR validation confirmed that skin neutrophil significantly upregulates *Il17a* for ten-fold among IL-17 family members measured (Figure 2D). These results indicate that neutrophils in the inflamed skin might play a pathological role in promoting psoriatic skin inflammation via IL-17A expression, a key driver for psoriasis.^29^

**Figure 2.**
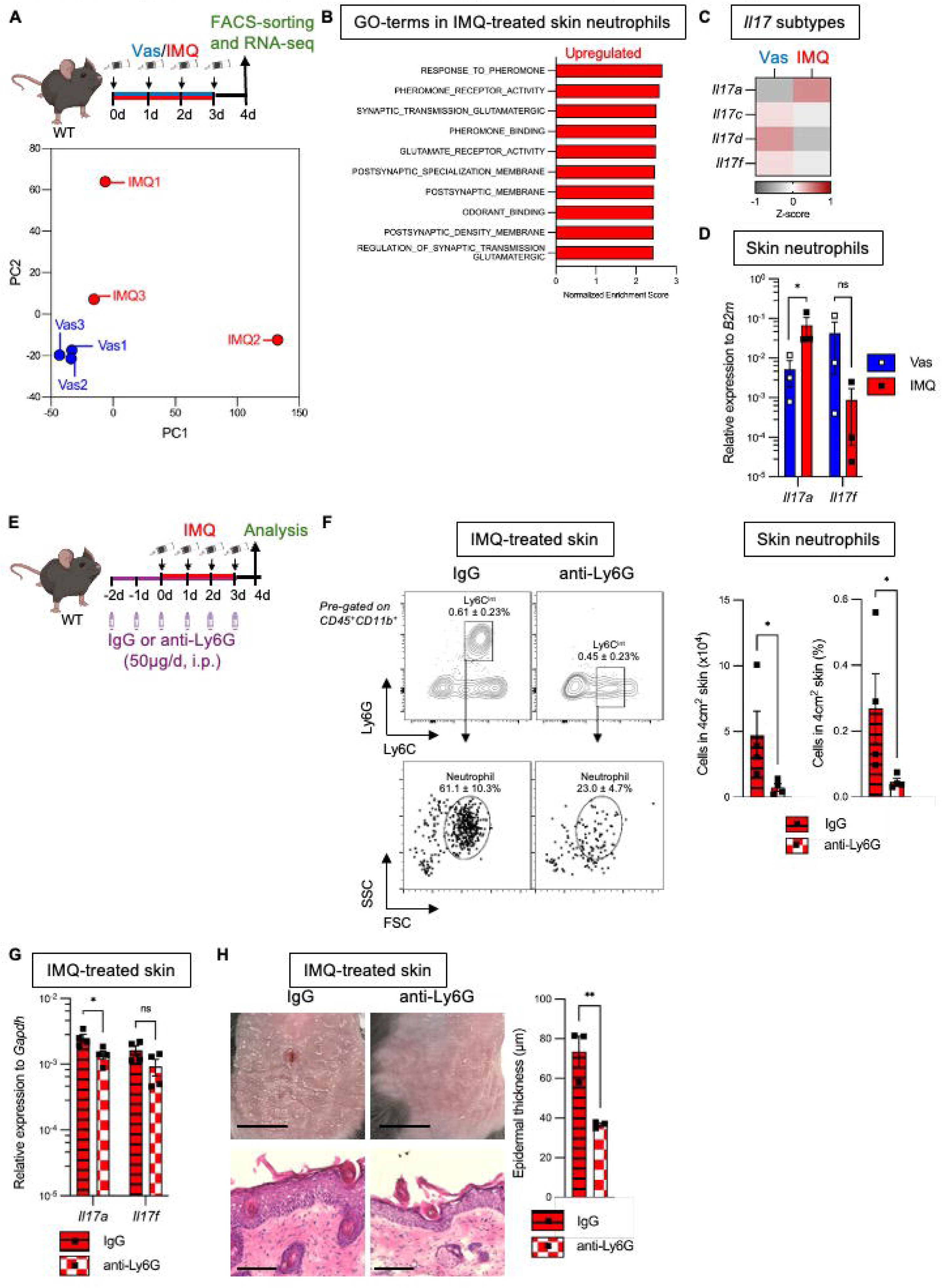
Skin neutrophils reinforce IL-17A signal and mediate psoriasis pathology. **(A)** Experimental scheme and PC plot of the skin-neutrophil bulk RNA-seq which results are shown in (B-D). **(B)** Gene ontology analysis of neutrophils from Vas-treated (n=3) and IMQ-treated skin (n=3). **(C)** Heatmap of IL-17 family gene expression in skin neutrophils after treatment with Vas (n=3) and IMQ (n=3), based on RNA-seq. **(D)** RT-PCR of *Il17a and Il17f* subtypes from sorted skin-neutrophils treated with Vas (n=3) and IMQ (n=3). **(E)** Experimental scheme of *in vivo* neutrophil depletion with anti-Ly6G neutralizing and IgG isotype-matched control antibodies which results are depicted in (F-H). Antibodies were injected once a day starting at 2d before topical IMQ-treatment (-2d) to 3d (total 6x injections). Analysis was conducted at 4d. **(F)** Flowcytometric analysis of skin neutrophils from IMQ-skin injected with neutralizing antibodies. Representative plot pre-gated on CD45^+^CD11b^+^ followed by the bar chart quantifying the absolute cell number and proportion of skin-neutrophils (IgG: n=4; anti-Ly6G: n=4). **(G)** RT-PCR results of *Il17a and Il17f* in the IMQ-skin injected with IgG (n=4) and anti-Ly6G (n=4). **(H)** Psoriasis clinical examination performed at the dorsal site. The dorsal skin images (upper panel: naked skin with bar indicating 1cm; lower panel: histology with black line reflecting 100µm). Epidermal thickness of the dorsal IMQ-skin intraperitoneally administered with IgG: n=3; anti-Ly6G: n=3). Data are pooled from ≥ 2 independent experiments with each dot shown in the chart represents the measurement from 1 experimental subject and shown as mean±S.E. \**p*<0.05; \*\**p*<0.01.

To better characterize the function of neutrophil in psoriasis pathology, we depleted the neutrophil by daily injection of anti-Ly6G neutralizing antibody while applying IMQ over 6 days (Figure 2E). Because antibody binding masks Ly6G for immunophenotyping, neutrophils were quantified using size- and complexity-based gating (Figure 2F). In line with the previous study,^30^ this approach could achieve around 80% depletion in skin (Figure 2F) and about 60% reduction in the circulating neutrophils (Figure S2B). Neutrophil depletion markedly reduced *Il17a* expression in psoriatic skin (Figure 2G) and alleviated clinical (scaling, erythema) and histological (epidermal thickening) manifestations by approximately 50% (Figure 2H and S2A).

Altogether, these results suggest that psoriatic neutrophils are an important source of IL-17A and contribute to disease progression.

### Skin inflammation in psoriasis drives emergency granulopoiesis in bone marrow

Neutrophils are short-lived innate immune cells that rapidly respond to tissue insults and initiate downstream immune reponses. Upon antigenic clearance, neutrophil demand rises sharply and met through emergency granulopoiesis in the BM.^8,31^ Given that psoriatic skin is infiltrated by a large number of neutrophils (Figure 1A-D), we reasoned that BM hematopoiesis might be activated to meet the demand. To test this, we first examined early hematopoietic compartment in BM during psoriasis (Figure 3A), including hematopoietic stem and progenitor cell (HSPC) fractions such as Lin^-^ c-Kit^+^Sca-1^+^ (LSK) or Lin^-^c-Kit^+^CD86^+^ (L86K), an alternative HSPC immunophenotyping under inflammatory conditions.^32^ Generally, LSK-defined HSPCs expanded as early as 16h post-psoriasis induction (Figure S3A-S3C). LSK CD150^+^CD48^-^ long term HSC (lt-HSC) increased toward 4d, whereas LSK CD150^-^CD48^-^ short term HSC (st-HSC) rapidly increased at 16h and decreased thereafter. In contrast, the myeloid-biased multipotent progenitor (MPP2/MPP3)^33^ but not lymphoid-biased MPP4 rapidly expanded by 5-to 9-fold at 16h post IMQ treatment, compared to control Vas, and remained elevated during the disease course (Figure 3B). Similarly, L86K-defined HSPC profiling showed an early increment of myeloid-biased MPP2/MPP3 reaching more than 5-fold increase upon psoriasis, with an overall reduced HSC fraction and unaltered MPP4 number (Figure S4B and S4C). Both immunoprofilings consistently suggest that the early hematopoiesis in the BM rapidly responds to distal skin inflammation and is skewed toward myelopoiesis, similar to the other systemic inflammation.^34^

**Figure 3.**
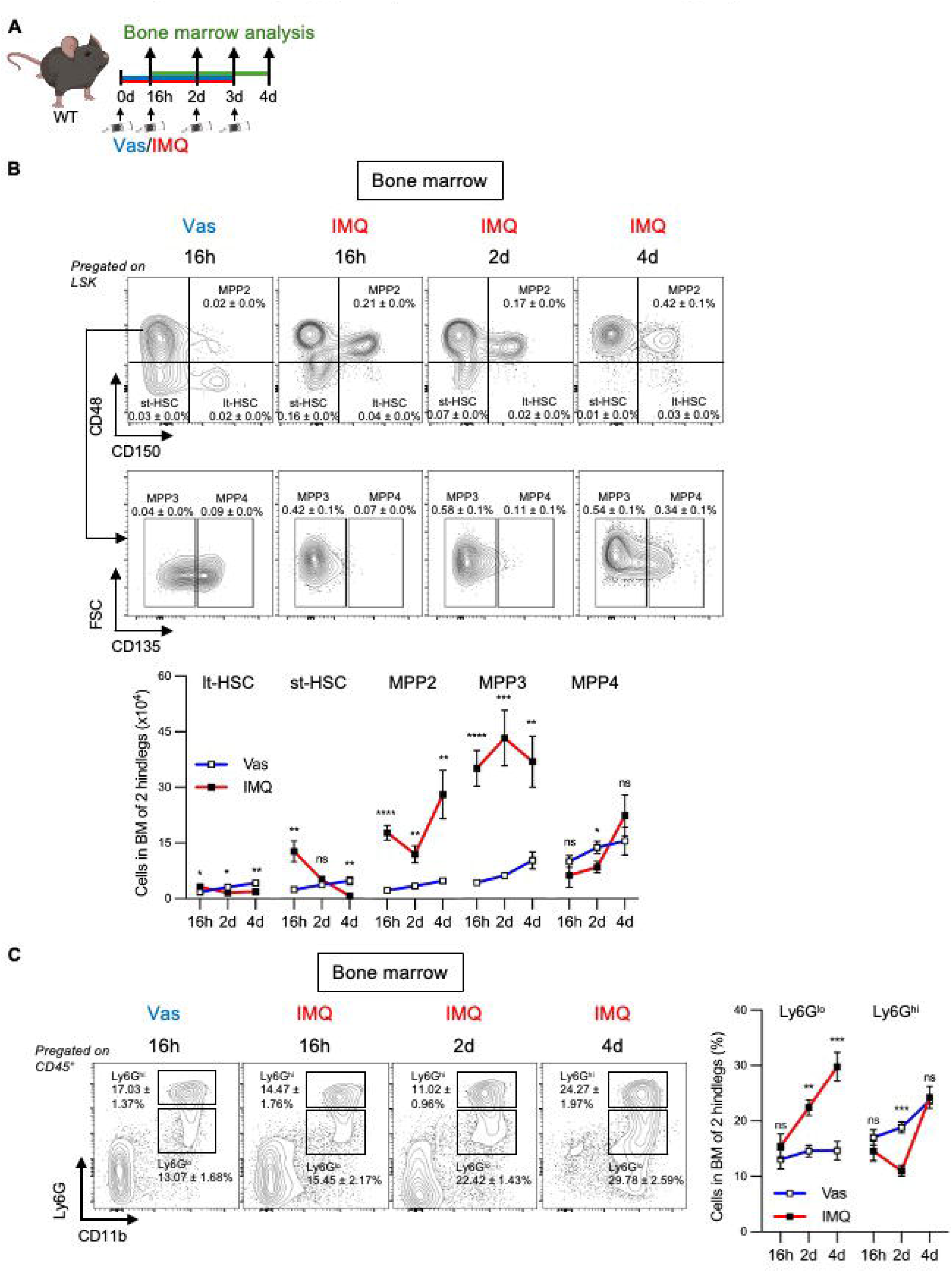
Emergency granulopoiesis activated in the psoriatic bone marrow. **(A)** Experimental scheme of BM analysis. Mice were topically treated with Vaseline (Vas) or Imiquimod (IMQ) at the dorsal skin site once/day at 0-3d and analysis was done sequentially from different animals at 16h, 2d and 4d post-treatment. **(B)** FACS analysis of HSPC in the BM including the representative plots pregated on LSK cells in a time-course manner and the corresponding line plots depicting the absolute number of lt-HSC, st-HSC, MPP2, MPP3, and MPP4 in the BM after treatment with topical Vas (n=3-6) or IMQ (n=3-6). **(C)** FACS analysis of immature (Ly6G^lo^) and mature (Ly6G^hi^) granulocyte in the BM. Representative plots pregated on CD45^+^ BM cells (left) and time-course kinetics proportion (right) of Ly6G^lo^ and Ly6G^hi^ BM cells after topical Vas (n=3-5) or IMQ (n=3-5) treatment. Data are pooled from ≥2 independent experiments with each dot shown in the chart represents the measurement fro 1 experimental subject and shown as mean±S.E. \**p*<0.05; \*\**p*<0.01; \*\*\**p*<0.001; \*\*\*\**p*<0.0001.

We next analyzed granulopoiesis in BM which *de novo* generates neutrophils. It was found that psoriasis induced a 2-3 fold expansion of Ly6G^lo^ immature granulocytes within 4d, whereas Ly6G^hi^ mature granulocytes decreased by about 40% at 2d and returned to baseline by 4d (Figure 3C). In the steady-state skin, myeloid population accounts for less than half of CD45^+^ cells, with only about 1% neutrophil residing (Figure S3D). At 16h post psoriasis induction neutrohils reached 4%, further expanded to 8% at 2d, and 28% of the total hematopoietic (CD45^+^) compartment by 4d. Concurrently, CD11b^+^ granulocytes in the BM progressively expanded comprising immature fraction from 13% to 30% and mature granulocytes from 17% to 24% of total BM cells at 4d after psoriasis induction. The synchronous expansion of BM and skin granulocytes (Figure S3D) indicates a central role of BM to generate and directionally efflux neutrophils to skin upon psoriasis.

Taken together, these findings demonstrate that psoriatic skin inflammation activates emergency granulopoiesis in the BM, enforcing the cutaneous neutrophil burden in the inflammed skin.

### Endothelial cells in piratic skin produces G-CSF leading to its systemic elevation

Emergency granulopoiesis is driven by myeloid-supporting cytokines.^8,35^ To identify the mediators of skin-BM crosstalk during psoriasis, we tested various proinflammatory cytokines in the serum of psoriatic mice (Figure 4A and S4A). Among those tested (data not shown), IL-1α increased by approximately 3-fold whereas IL-1β decreased by half. Tumor necrosis factor-α (TNF-α) and GM-CSF remained unchanged (Figure S4B). Since IL-1α is known to induce granulopoiesis,^36^ we blocked IL-1 signaling by daily injection of an IL-1 receptor (IL-1r) antagonist, Anakinra over 4 consecutive days (Figure S4C).^37,38^ The IL-1r blockade did not significantly affect neutrophil expansion in BM and skin (Figure S4D-E), nor psoriasis pathology including gross (scaling and erythema) and histological (epidermal thickness) manifestations (Figure S4F). These results indicate that IL-1 signaling does not play an important role for psoriasis-associated skin-BM crosstalk.

**Figure 4.**
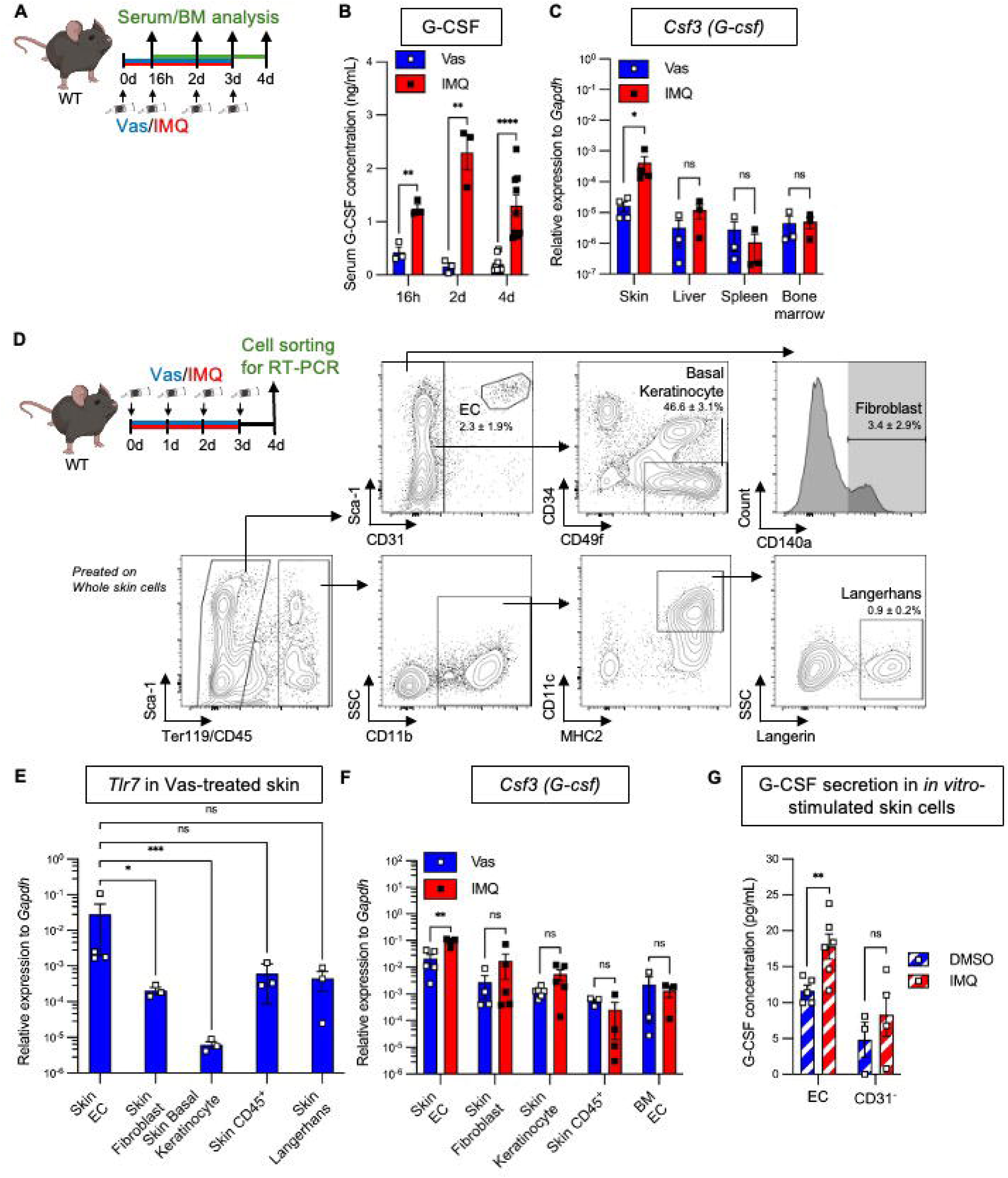
Psoriasis induces G-CSF expression in skin endothelial cells. **(A)** Experimental scheme of serum and BM analysis for results depicted in (B and C): Mice were treated with Vas or IMQ once/day at the dorsal skin at 0-3d and analysis were done successively at 16h, 2d and 4d with different mice from each time-point. **(B)** Time-course kinetics of serum G-CSF concentration after treatment with Vas (n=3-9) or IMQ (n=3-9). **(C)** Relative expression of *Csf3 (G-csf)* mRNA in skin, liver, spleen, and bone marrow at 4d post-treatment with Vas (n=3-4) or IMQ (n=3-4). **(D)** Experimental scheme and the representative gating strategy for skin cell subtype by flowcytometric sorting at 4d after topical Vas/IMQ treatment which results are depicted in (E and F): a hundred cells were sorted for each subtype (endothelial cells (ECs), basal keratinocyte, fibroblast, Langerhans cells). **(E and F)** Relative expression of *Tlr7* **(E)** and *G-csf* **(F)** mRNA (Vas: n=3-5, IMQ: n=3-5) from the cellular subsets mentioned in the respective bar graphs. **(G)** Sixty thousand cells were sorted and planted into culture media with or without IMQ (50µg/mL) for 48h and G-CSF was measured from the media subsequently (n=3-6). Data are pooled from ≥2 independent experiments with each dot shown in the chart represents the measurement from 1 experimental subject and shown as mean±S.E. \**p*<0.05; \*\**p*<0.01; \*\*\**p*<0.001; \*\*\*\**p*<0.0001.

We next examined G-CSF, a key driver of granulopoiesis (Figure 4A). G-CSF protein level in the serum rose sharply, about 5-fold at 16h and up to15-fold by 2d to 4d post psoriasis induction, concurrent with the upstream HSPC activation (Figure 4B, 3B and S3A). To identify which organ produces G-CSF, we compared gene expression *Csf3* (*G-csf*) in skin, liver, spleen, and BM at 4d,^9^ and found that only skin showed a significant upregulation of *Csf3* upon psoriasis (Figure 4C).

To identify what skin cell subtypes contribute to psoriasis-induced G-CSF secretion, we flow cytometrically sorted specific skin cell populations including endothelial cells (ECs), fibroblasts, basal keratinocytes, CD45^+^ hematopoietic cells, and langerhans cells and compared their gene expression (Figure 4D). Expression of *Tlr7*, the IMQ receptor was highest in skin ECs and >100-fold higher than other subsets in steady-state skin (Figure 4E). *Csf3* was significantly higher in steady-state skin ECs than other cells tested (*p*<0.05) and significantly upregulated only in skin ECs upon psoriasis (Figure 4F). Of note, the BM-ECs showed no *Csf3* change even after psoriasis induction, indicative of skin EC-specific G-CSF production induced by psoriasis (Figure 4F).

To directly test the capacity of skin ECs to secrete G-CSF in response to IMQ treatment, ECs and CD31^-^ cells were sorted from steady-state skin and cultured in the presence or absence of IMQ for 2 days. In the absence of IMQ, G-CSF secretion by cultured skin ECs was more than twice of skin CD31^-^ cells. Upon IMQ stimulation, skin ECs increased G-CSF secretion by approximately 50% over baseline, whereas skin CD31^-^ cells did not show any increase at all (Figure 4G). These results suggest that skin-resident ECs are a major producer of G-CSF in response to TLR7 stimulation.

Taken together, skin-resident ECs are able to respond to a TLR7 agonist via its receptor expression and readily secrete G-CSF upon stimulation, possibly contributing to its systemic elevation during psoriasis.

### Blockade of emergency granulopoiesis mitigates psoriasis

To evaluate the significant contribution of skin-derived G-CSF in psoriasis pathology, we systemically administered anti-G-CSF neutralizing antibody *in vivo*^39^ before and during psoriasis induction (Figure 5A). Anti-G-CSF treatment reduced circulating neutrophils in psoriatic mice by approximately 80% at 4d post induction (Figure S5A). BM analysis revealed that the treatment led to 30% suppression of psoriasis-induced LSK cell expansion in BM (Figure S5B). Blocking G-CSF specifically diminished the myeloid-biased MPP3 expansion by about 60%, while no significant effect on HSCs and other MPPs were observed (Figure 5B). Its inhibition resulted in a significant reduction (ca. 30%) of psoriasis-induced granulocyte expansion for both Ly6G^lo^ immature and Ly6G^hi^ mature granulocytes (Figure 5C). Consistent with suppressed BM granulopoiesis, neutrophil burden in skin decreased by >80%, accompanied by reduced clinical (erythema, scaling) and histological (epidermal thickening) severity by about 50% (Figure 5E and S5C).

**Figure 5.**
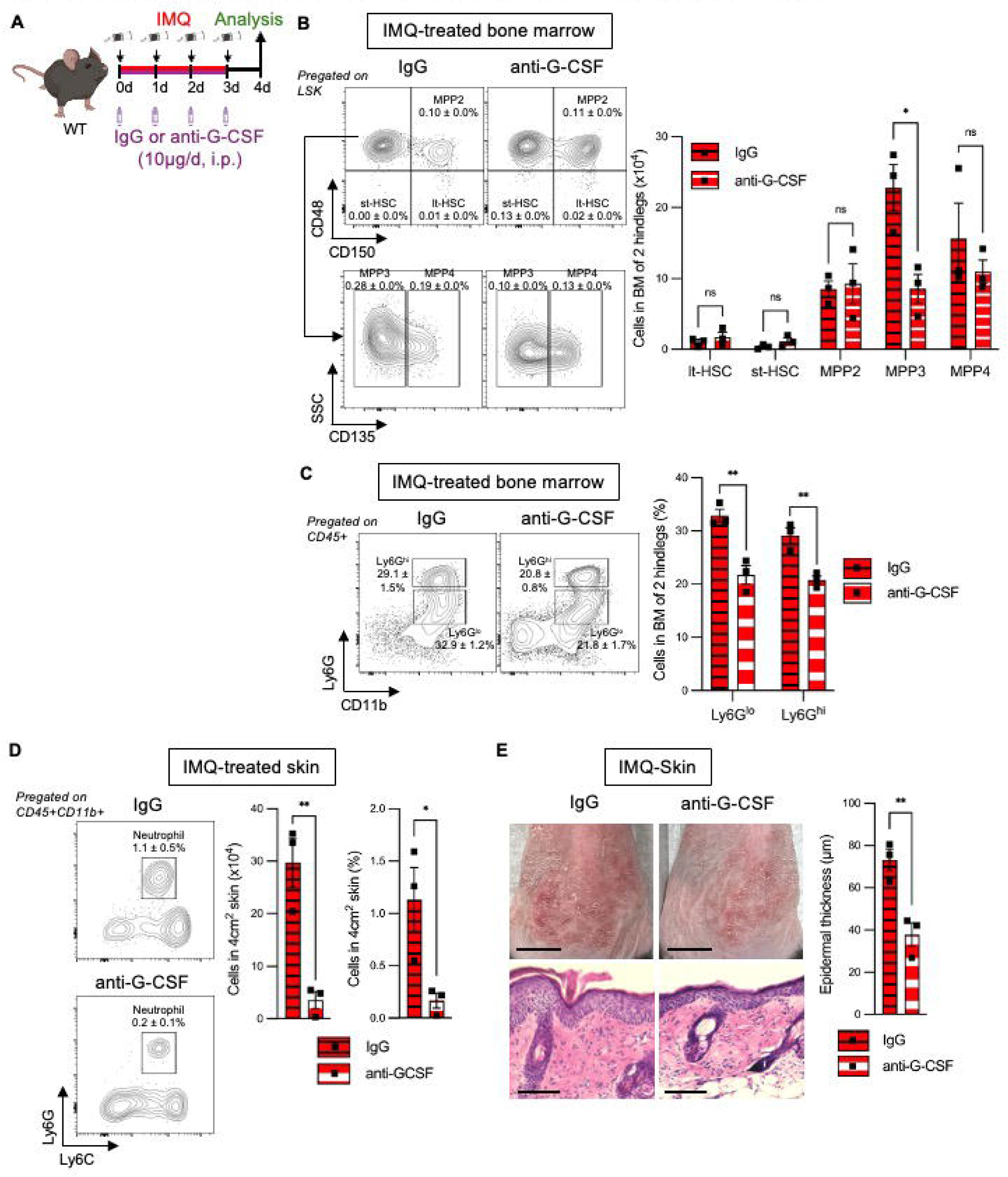
Inhibition of G-CSF-mediated granulopoiesis alleviates psoriasis symptoms. **(A)** Experimental scheme of G-CSF neutralization (B-E): mice were i.p. pre-injected with anti-G-CSF neutralizing or IgG isotype-matched control antibodies at 30 minutes before daily topical-IMQ application for 4 consecutive days (0-3d). Analysis was performed at 4d post-treatment. **(B)** FACS analysis of the BM showing representative FACS plot (left) and bar graphs (right) of absolute cell numbers of phenotypic lt-HSC, st-HSC, MPP2, MPP3 and MPP4 cells pre-gated on BM LKS co-treated with IMQ and with IgG (n=3) or anti-G-CSF (n=3). **(C)** FACS analysis of immature (Ly6G^lo^) and mature (Ly6G^hi^) granulocytes pregated on CD45^+^ BM cells. Representative FACS plot (left) and proportion (right) of both immature (Ly6G^lo^) and mature (Ly6G^hi^) granulocyte fractions from IMQ-BM treated with IgG (n=3) or anti-G-CSF (n=3). **(D)** FACS analysis of skin-resident neutrophils pre-gated on CD45^+^CD11b^+^ cells from dorsal skin. Representative FACS plot (left), number (middle) and proportion (right) of neutrophils in the skin co-treated with IMQ and IgG (n=3) or anti-G-CSF (n=3). **(E)** Representative naked skin images with scale bar of 1cm (left upper panel) and skin histology with HE staining (lower panel, black line indicates 100µm) (left lower panel). Quantification of epidermal thickness in skin co-treated with IMQ and IgG (n=3) or anti-G-CSF (n=3) (right). Data are pooled from ≥2 independent experiments with each dot shown in the bar chart represents the measurement from 1 experimental subject and shown as mean±S.E. \**p*<0.05; \*\**p*<0.01.

Collectively, our data suggest that G-CSF is a critical factor in orchestrating pathological skin-BM coordination of cutaneous neutrophilic inflammation exemplified in psoriasis.

### Neutrophil activation and G-CSF expression in endothelial cells of human psoriatic skin

To assess whether our findings in the mouse model is relevant to human psoriasis, we reanalyzed public bulk RNA-seq dataset (GSE54456) containing nearly 100 skin biopsies of normal and psoriatic patients.^19^ The dataset showed clear segregation of normal and psoriatic samples, with a significant upregulation of hallmark dermatitis cytokines including around 3-log increase in *IL17A, IL17F, IL23,* and *IL22, and* 2-log increase in *TNFA* and *IL6* in the psoriatic skin biopsies (Figure S6A and S6B).^1,26^ The psoriatic skin showed upregulated expression of neutrophil-recruiting chemokines such as *CXCL1, CXCL2, and CXCL5* (≥10 time for all),^40^ along with 5-time increase in *CCL2* (Figure 6A). Moreover, a significant elevation of neutrophil activity-related genes was observed, such as activation molecules (*S100A8, S100A9*) by about 100-fold, oxidative activity *(MPO*, *NOX2*) by about 2-fold (Figure 6A). An increase of phagocytic initiator *FCGR2A/FCGR3B* were observed, whereas serine protease *ELANE* and *CSTG* were found downregulated, suggesting an oxidative non-proteolytic phagocytosis action. Consistent with the findings in murine model, granulopoiesis-driving factors such as *CSF2* (*GM-CSF*) *and CSF3* (*G-CSF*) were elevated about 10-fold in psoriatic skin (Figure 6A).^41^ These results suggest an active recruitment and enhanced activity of neutrophil as well as increased level of neutrophil-supporting cytokine in human psoriatic skin.

**Figure 6.**
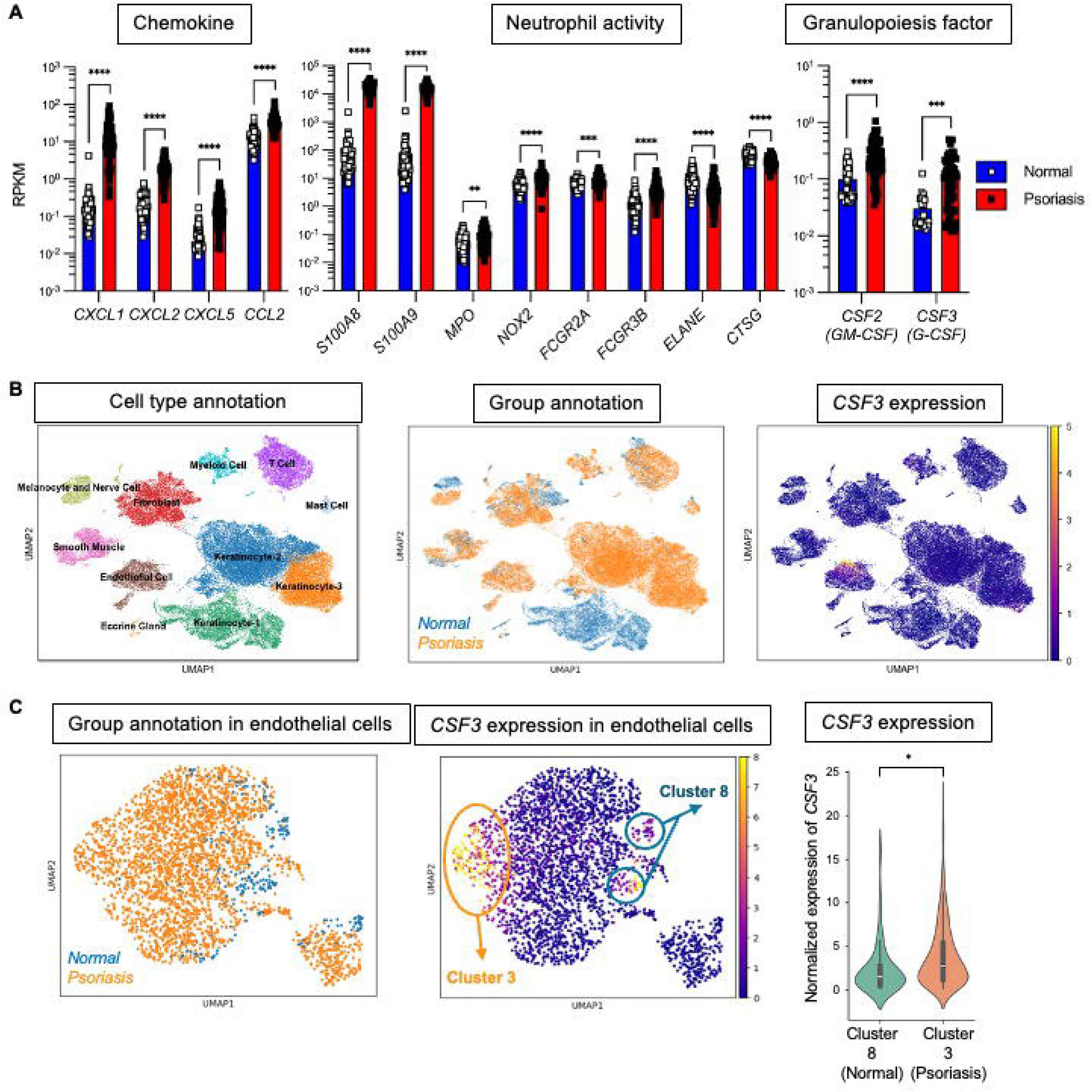
Human psoriatic skin shows neutrophil overactivity and elevated G-CSF expression in endothelial cells. **(A)** Expression of genes related to neutrophil function in the publicly available human psoriasis database (GSE54456). Each dot represents individual skin biopsy samples (normal: n=82, psoriatic skin: n=92). Expression of mRNA related to chemoattractant (left), neutrophil activity (middle), and granulopoiesis factor (right) (*CSF*: colony stimulating factor, *CSTG*: cathepsin-G, *CXCL*: CXC chemokine ligand, *ELANE*: neutrophil elastase, *FCSGR2A: Fc*γR*-2a, FCSGR2A: Fc*γR*-*3b*, IL*: interleukin, *MPO*: myeloperoxidase, *TNF*: tumor necrosis factor). **(B)** Clustering analysis of single cell RNA-seq dataset (GSE173706) of the skin biopsies (normal: n=8, soriasis: n=14) covering the annotation of the cell type clusters (left), annotation of the cluster’s disease condition (normal/psoriasis) (middle), and the differential expression of *CSF3* mRNA among all the annotated clusters (right). **(C)** Endothelial cell subsetting and re-analysis (GSE173706), covering group annotation of the in the endothelial cell subsets’ disease condition (normal/psoriasis) (left), differential expression analysis of *CSF3* (middle), and normalized *CSF3* expression of re-analyzed endothelial cells from cluster 8 (normal) and cluster 3 (psoriasis) (right). Data are shown as mean±S.E. \**p*<0.05; \*\**p*<0.01; \*\*\**p*<0.001; \*\*\*\**p*<0.0001.

To explore the underlying cellular mechanisms, we looked into a single-cell RNA-seq dataset (GSE173706) and annotated 11 distinct clusters including keratinocyte type 1-3, melanocyte, nerve, eccrine gland, endothelial cell, fibroblast, smooth muscle, T cell, myeloid cell, and mast cell (Figure 6B, S6C and S6D).^20^ The keratinocyte type 1 belongs to normal skin and is differentiated from type 2 and 3 found in psoriatic skin, a hallmark change in psoriatic skin previously reported (Figure 6B, S6E).^42^ Human psoriatic skin showed about double numerical expansion of both EC and myeloid cell compartments. Notably, *CSF3* expression was restricted to a subset of ECs (Figure 6C). Re-clustering of ECs revealed two distinct *CSF3* positive subsets: Cluster 8 of normal skin and Cluster 3 of psoriatic skin (Figure 6C, S6F). The *CSF3* expression was significantly higher in psoriatic ECs (Cluster 3) compared to normal ECs (Cluster 8) (Figure 6C).

Taken together, our findings suggest that human psoriatic skin is found to contain activated neutrophils that secrete psoriasis-mediating factors as well as endothelial cells that enhances G-CSF expression, similarly to a murine psoriasis.

## DISCUSSION

In this study, we demonstrate that (1) upon psoriasis, skin-resident endothelial cells are able to sense TLR7 agonist and secrete G-CSF, (2) which skews hematopoiesis in BM into emergency granulopoiesis, (3) concurrently, some neutrophils egress BM and progressively infiltrate into the inflamed skin through expanded blood vessels upon psoriasis, and (4) promotes tissue inflammation, such as IL-17A and worsening disease symptoms. (5) Depletion of neutrophils or blockade of G-CSF-mediated emergency granulopoiesis alleviates disease symptoms. (6) The similar mechanism might exist in human psoriasis.

Endothelial cells composing of the vascular network, not only regulate leukocyte trafficking but also actively interact with immune cells to regulate their function.^43^ ECs have been shown to secrete GM-CSF and hence stimulating neutrophil degranulation *in vitro*,^44^ whereas BM-resident ECs increase G-CSF production in response to systemic pathogen signals *in vivo*, promoting neutrophil generation and egress.^9,45^ Our findings support that in psoriasis, skin-resident ECs act as the key regulator of cross-organ communication by driving BM granulopoiesis through G-CSF production. This cross-organ communication is evidenced by other studies showing that stress in epithelial barrier can promote systemic myelopoiesis in various organs.^46,47^ Our data reveals that upstream myeloid progenitors respond to G-CSF and expand more quickly than the downstream immature neutrophils (Figure 3 and S3), in accordance with expression level of G-CSF receptor.^48^ Given that G-CSF also serves as a critical neutrophil survival factor,^49,50^ the close proximity of neutrophils to skin ECs within the psoriatic lesion would likely help to prolong their lifespan and to populate them in the inflamed skin. Collectively, these findings highlight a pathologic role of skin-resident ECs as a niche to promote infiltrating neutrophil and drive psoriasis pathology remotely and locally by sending granulopoietic and survival G-CSF signals, respectively.

During inflammation, tissue neutrophils typically enhance phagocytic and metabolic activity to promote reactive oxygen species/protease production and NETs for immunogen clearance.^51–53^ However, our RNA-seq data did not show clear evidence that neutrophils in psoriatic skin, upregulate these functions, but instead that they did increase response to neurochemical signals. The skin microenvironment is a multi-compartmental organ tightly controlled by innervated neuron,^54,55^ where neutrophils express neurotransmitter receptors which regulate their chemotaxis,^56,57^ in addition to chemokine-driven tissue recruitment.^40^ Notably, psoriatic neutrophils upregulate *Il17a,* and anti-Ly6G antibody-mediated depletion of neutrophil results in reduction of *Il17a* in skin tissue and mitigation of psoriasis (Figure 2G and 2H), demonstrating their key role in IL-17A-driven inflammation and confirming prior reports on IL-17A-expressing neutrophils in psoriatic skin.^27,28^

Among granulopoiesis-supporting factors, G-CSF and GM-CSF have overlapping but distinct effect on hematopoietic lineage instruction: GM-CSF broadly expands diverse myeloid cell types, whereas G-CSF selectively drives robust neutropoiesis.^58^ Anti-GM-CSF clinical trials in psoriasis reduced myeloid cell number but shows limited efficacy,^59,60^ whereas G-CSF supplementation increased neutrophil count which could trigger psoriasis dermatitis adverse events, ^61–64^ suggesting a pathogenic role of neutrophil in the disease progression consistent with our findings. Downstream of G-CSF signaling, JAK-STAT inhibitors show clinical promise, achieving sustained reduction up to 75% for as long as 40 weeks.^65,66^ Moreover, anti-TLR7 therapy for psoriasis has shown encouraging results and been advanced to clinical trials.^67–69^ Our findings add a mechanistic rationale for these observations by demonstrating that TLR7 sensing in skin ECs induces G-CSF secretion, driving BM-emergency granulopoiesis, leading to cutaneous neutrophil accumulation that aggravates psoriasis.

While many studies uncovered local cellular interactions in psoriatic skin, e.g. involving fibroblast, keratinocytes, and neutrophils,^70–72^ emerging evidences suggest that psoriatic skin inflammation pertains to a systemic consequences through cross-organ communication. A study demonstrated that transgenic mice with keratinocyte-specific IL-17A overexpression developed chronic psoriasis and bone loss due to osteoblast-osteocyte dysfunction.^73^ Moderate-to-severe psoriasis is also linked to myocardial infarction, likely driven by a combination of similar cytokines promoting both pathologies.^74,75^ Moreover, even acute psoriasis can cause increased susceptibility of developing gut colitis due to intestinal dysbiosis driven gut-macrophage hyperactivity.^76^ In our study, we identified a pathological cross-organ feedback loop in psoriasis initiated by insulted skin ECs. Although psoriatic skin ECs are known to be functionally altered and promoting local T-cells infiltration,^77^ we show that they also drive BM emergency granulopoiesis via G-CSF secretion, revealing a systemic role in skin-BM pathological crosstalk upon psoriasis.

Our findings suggest that level of circulating G-CSF or activity of skin-resident ECs could be used as biomarkers for disease severity and therapy response in psoriasis patients. It remains to be determined whether G-CSF driven skin-BM crosstalk is a distinctive feature of psoriasis or represents a broader neutrophil-associated skin inflammation such as atopic dermatitis or pustular dermatoses.^78,79^ Our study proposes the potential of targeting innate immunity as a complementary approach to existing biologics that primarily modulate adaptive immune pathways. Although IL-17 blockers (e.g. Secukinumab, Ixekizumab, Brodalumab) and IL-23 inhibitors (e.g. Guselkumab, Risankizumab, Tildrakizumab) show potential benefit, they provide limited control over neutrophil-driven inflammation.^80–83^ Given that neutrophil is key effector cells against infection, combining blockage of neutrophil infiltration to skin with adaptive immune blockers might therefore enhance therapeutic efficacy and potentially prolong remission.

## ACKNOWLEDGEMENTS

T.K. received a scholarship support from Ministry of Education, Culture, Sports, Science, and Technology-Japan (MEXT). We appreciate the logistical and technical assistance provided by International Core-facility of Advanced Life Science at Kumamoto University. We would also thank Dr. Tetsuro Kobayashi (RIKEN) for the help to establishing skin single cell suspension method. This work was supported by KAKENHI from JSPS (21KK0150, 21H02953 and 22K19548 to H.T.), KAKETSUKEN 2020 (The Chemo-Sero-Therapeutic Research Institute) (to H.T.), JST FOREST (JPMJFR200O to H.T.), The Japanese Society of Hematology (to H.T.), Joint research support at the Institute of Medical Science 2025 (to H.T.), The University of Tokyo, The Japan Prize Foundation 2023 (to H.T.). Takeda Science Foundation visionary research (to H.T.)

## AUTHOR CONTRIBUTIONS

T.K. designed and performed the experiments, analyzed the data, and wrote the manuscript. T.M. performed RNA-seq. S.L., J.Y., P.K., performed 3D-intravital imaging, K.W. and A.S. discussed and interpreted results. H.T. designed the experiments, supervised the research project, and wrote the manuscript. All authors read and approved the final manuscript.

## DATA AVAILABILITY STATEMENT

All data needed to examine the conclusions in this study are present in the paper and/or the Supplementary Materials. The RNA-seq data have been deposited in the DDBJ BioProject database (https://ddbj.nig.ac.jp/search) with accession number PRJDB37718.

## ETHICS STATEMENT

Animal experiments conducted in Kumamoto University and KAIST were performed under the approval of Animal Care and Use Committee of Kumamoto University and KAIST, respectively.

## CONTACT INFORMATION

Hitoshi Takizawa, PhD, Laboratory of Stem Cell Stress, International Research Center for Medical Sciences (IRCMS), Kumamoto University, Kumamoto 860-0811, Japan., Phone: +81 96 373 6847, Fax: +81 96 373 6869

