## Supplemental info for "Skin-derived G-CSF activates pathological granulopoiesis upon psoriasis"

Kosasih, *et* al. Figure S1. Characterization of clinical manifestations, myeloid cells, and the related factors in an imiquimod (IMQ)-induced psoriasis model

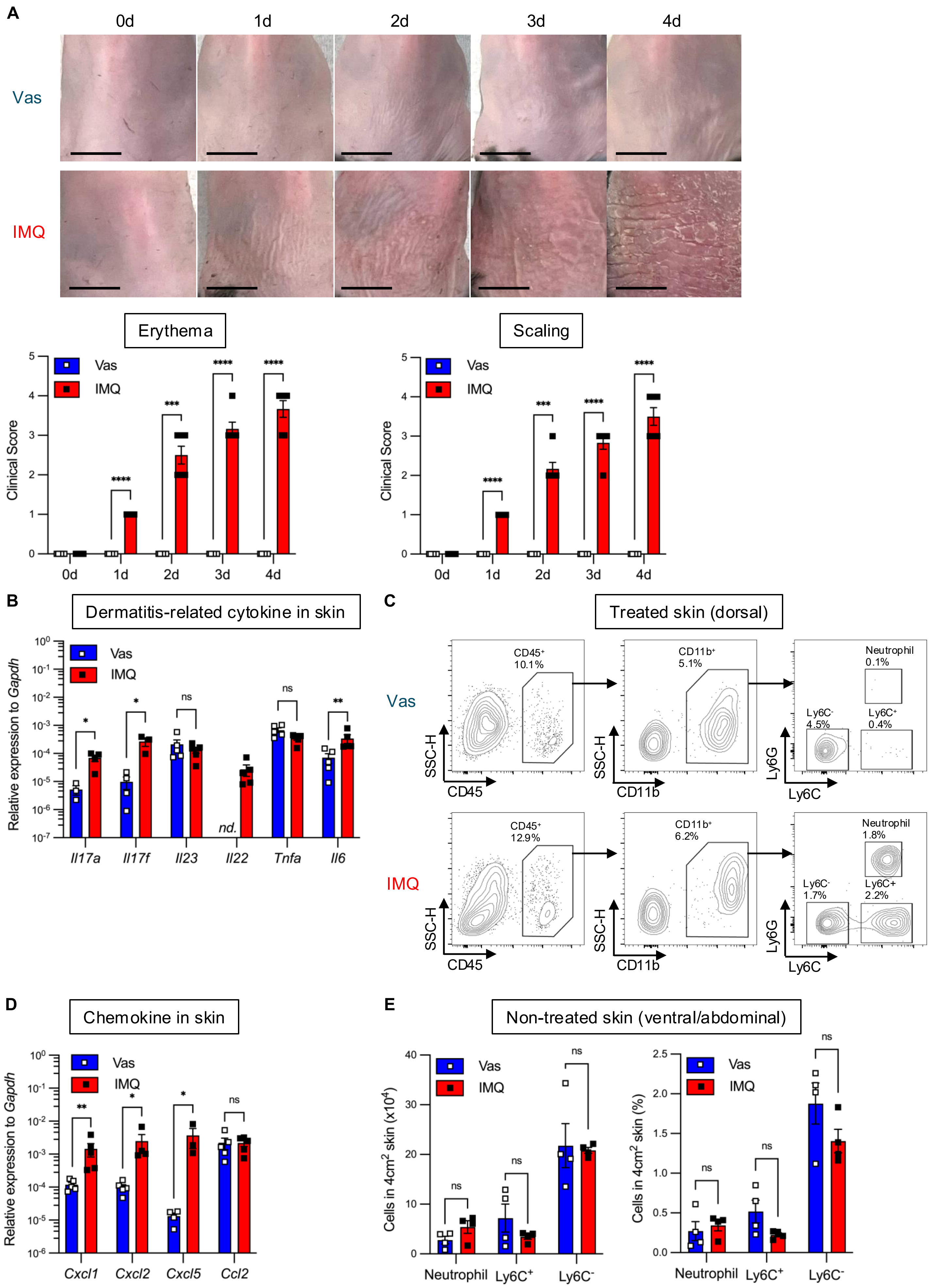

Kosasih, *et* al. Figure S1. Characterization of clinical manifestations, myeloid cells, and the related factors in an imiquimod (IMQ)-induced psoriasis model

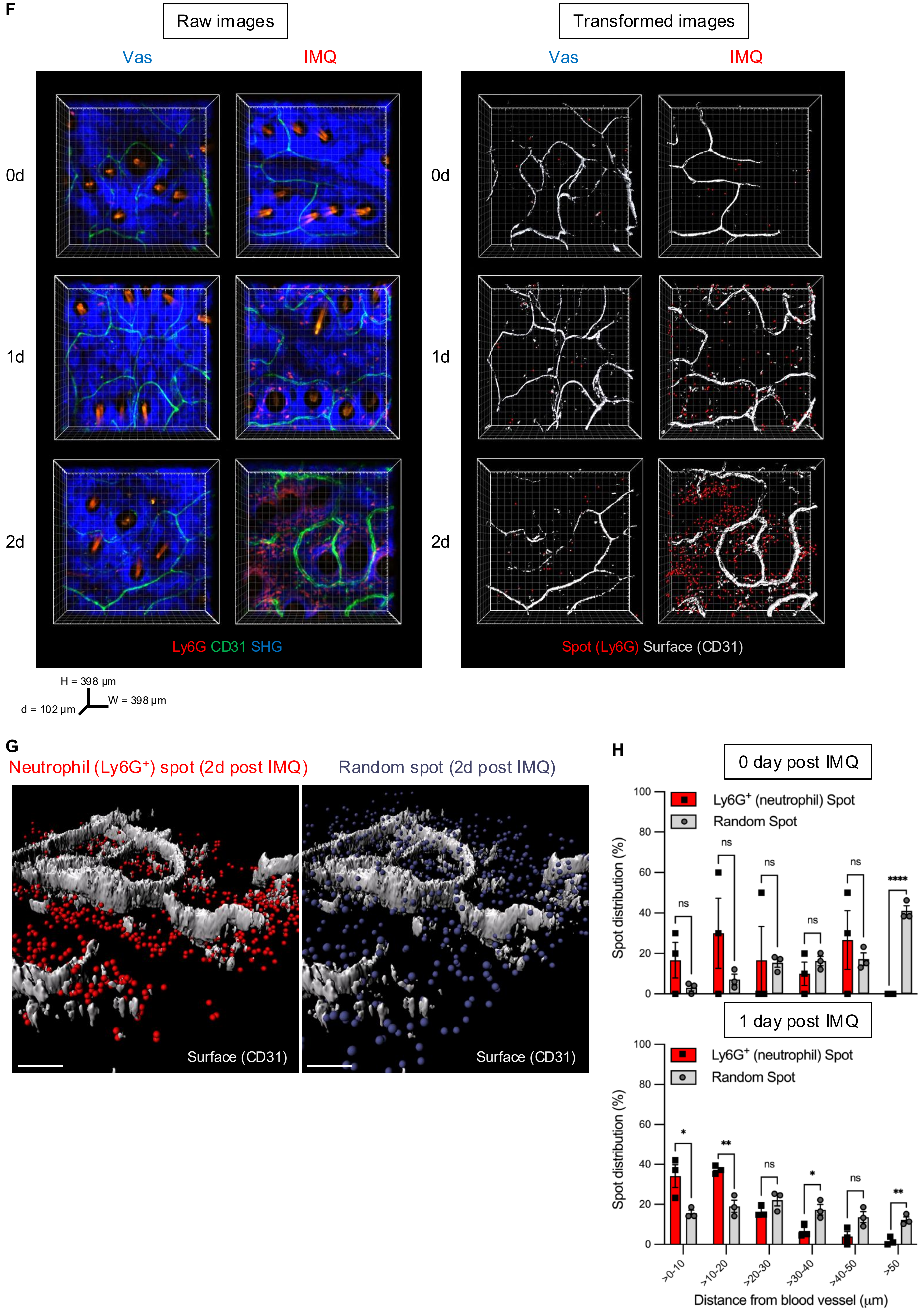

Kosasih, *et* al. Figure S2. Overactive neutrophils have pathological function in psoriasis.

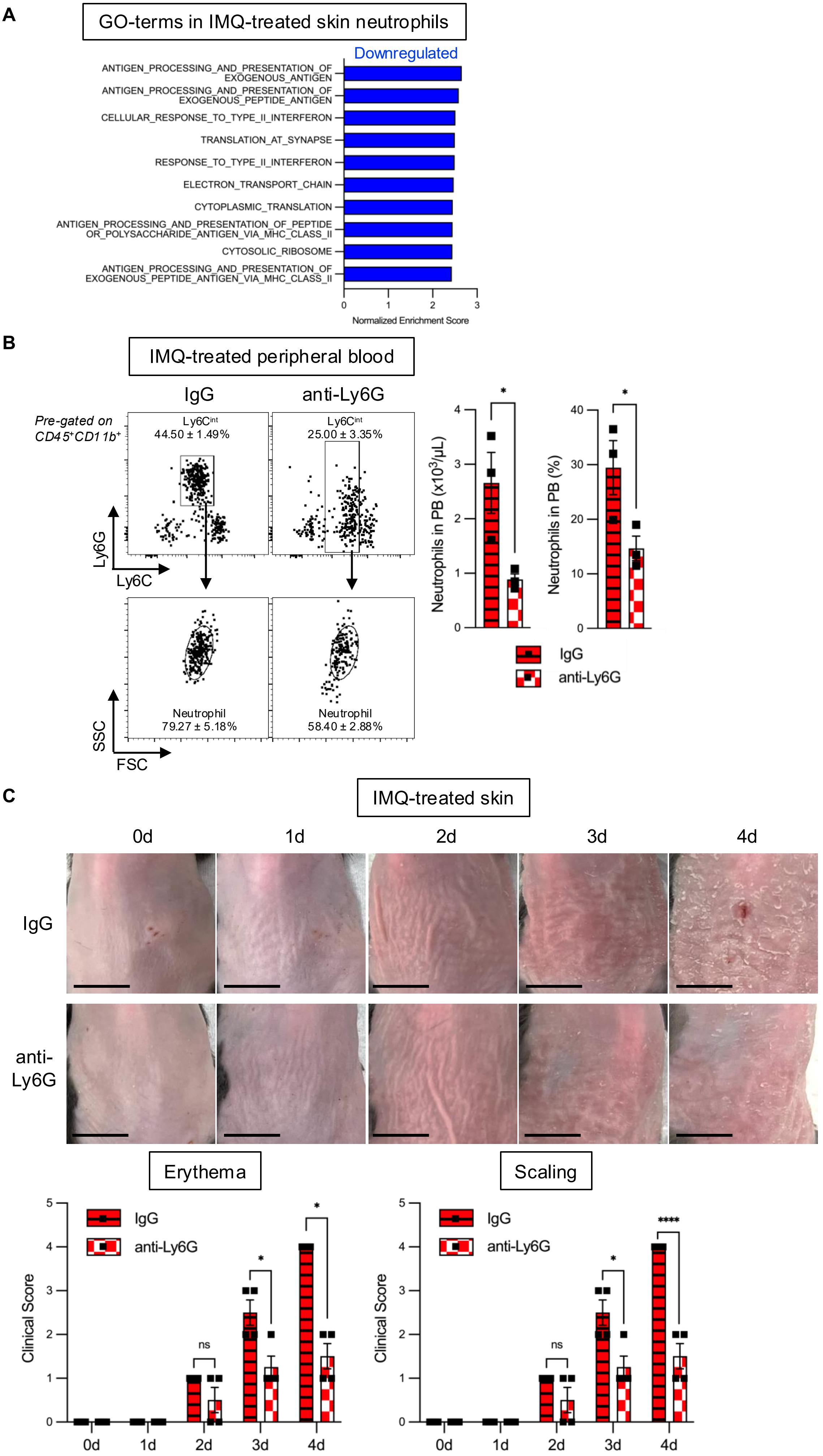

Kosasih, *et* al. Figure S3. Psoriasis induces expansion of L86K-defined HSPC in BM

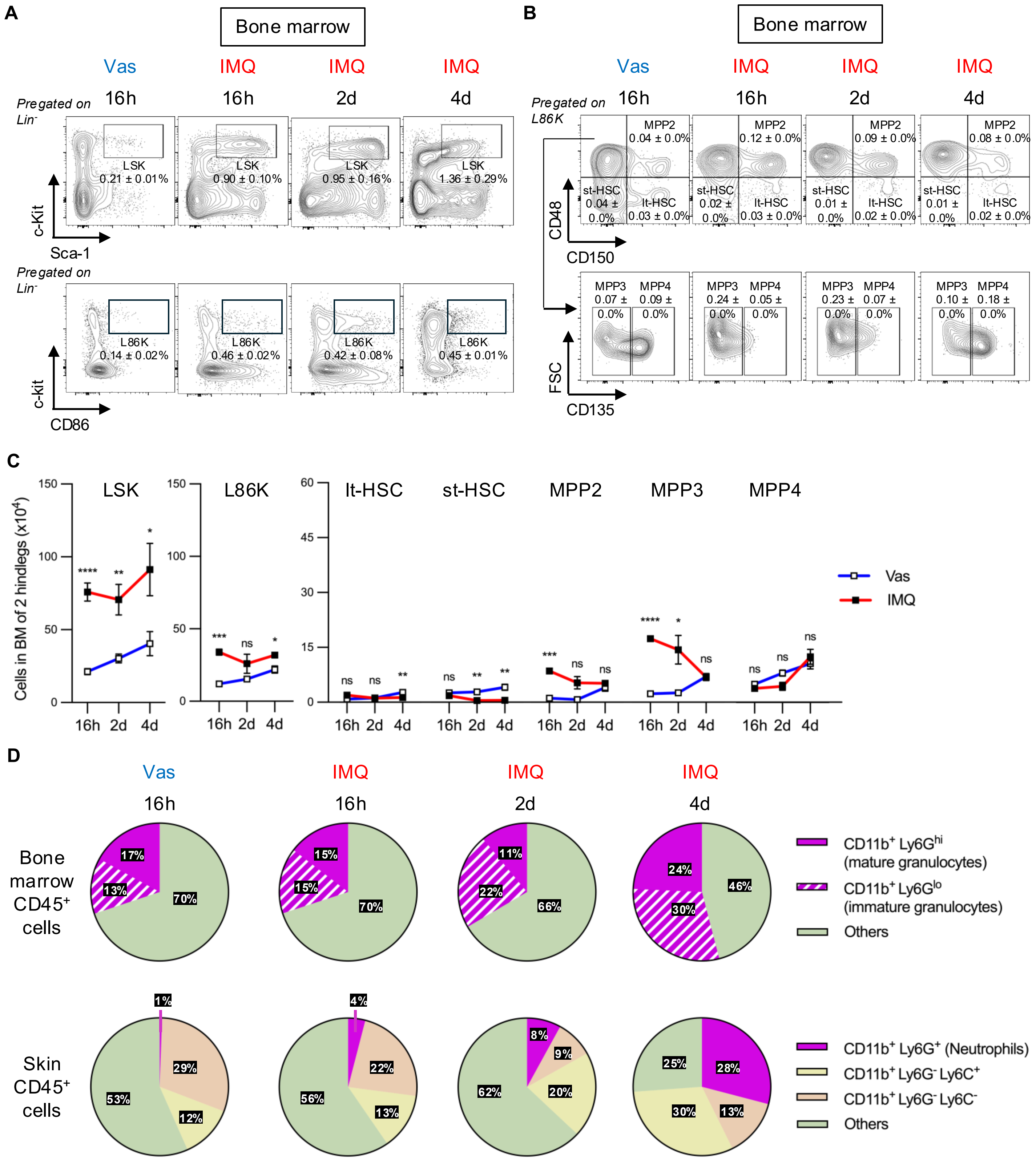

Kosasih, *et* al., Figure S4. IL-1 signals do not contribute to psoriasis-driven emergency granulopoiesis

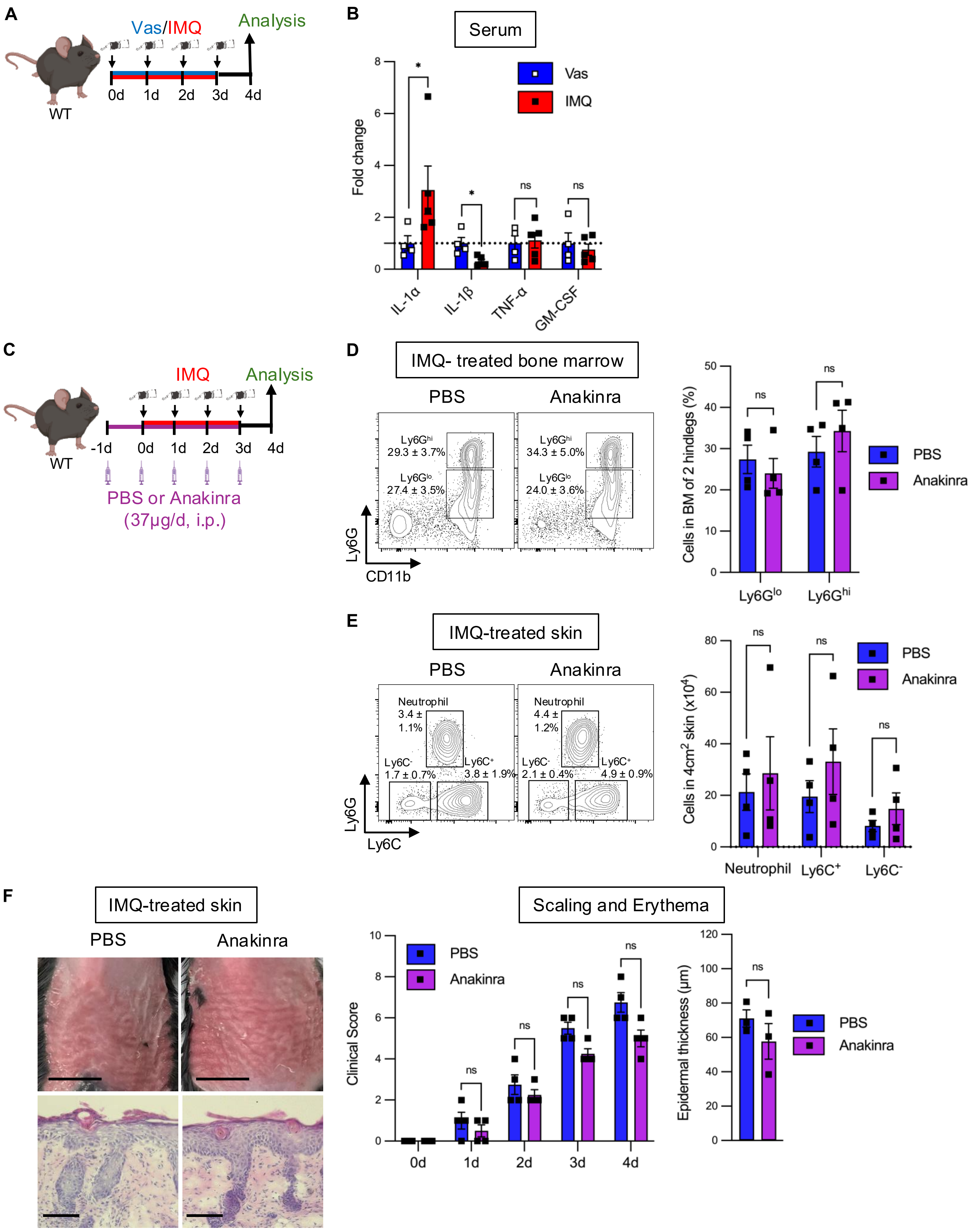

Kosasih, *et* al. Figure S5. G-CSF neutralization reduces circulating neutrophils, hematopoietic activation in bone marrow, and psoriasis clinical manifestation.

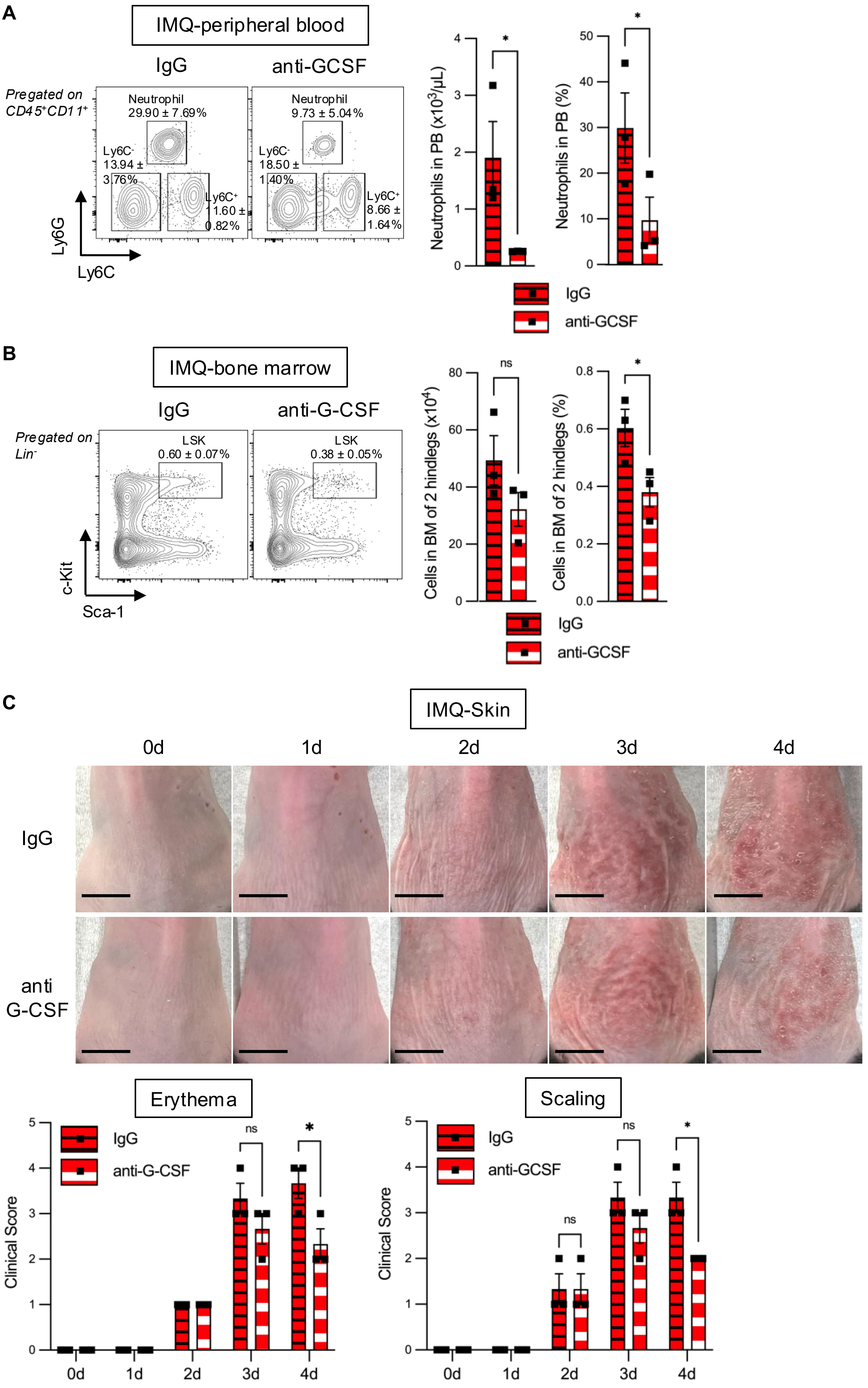

Kosasih, *et* al. Figure S6. Clustering and relative transcript measurement of human psoriasis skin biopsies

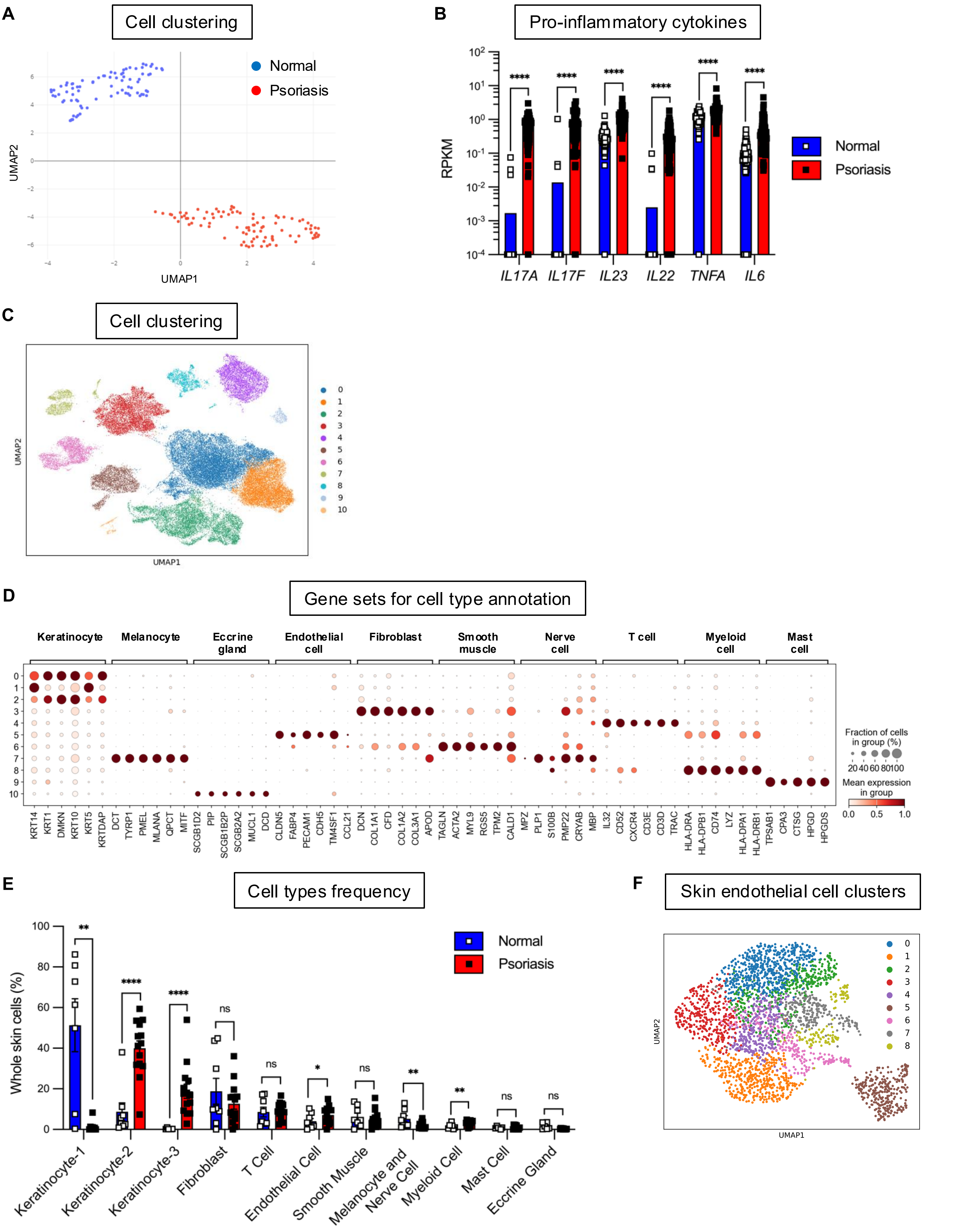

### SUPPLEMENTAL FIGURE LEGEND

**Figure S1 Characterization of clinical manifestations, myeloid cells, and the related factors in an imiquimod (IMQ)-induced skin psoriasis model.**

**(A)** Upper: representative images of dorsal skin treated topically with daily Vas/IMQ from 0-3d (4 times). Lower: Clinical score such as erythema and scaling from the dorsal skin treated with Vas/IMQ (n=3 each). **(B)** mRNA expression of dermatitis-related cytokines in dorsal skin treated with Vas (n=3-5) or IMQ (n=3-5 for each). **(C)** Representative FACS gating strategy plots of myeloid cell fractions from dorsal skin treated with Vas or IMQ. **(D)** mRNA expression of chemokines in the dorsal skin treated with Vas or IMQ (n=3-5 for each). **(E)** Number and percentage of myeloid cell fractions in the non-lesional abdominal skin (Vas: n=4, IMQ: n=4). **(F)** Representative 3-dimensional (3D) intravital microscopic images of dorsal skin at 0d, 1d, and 2d after Vas or IMQ treatment. Left images show raw data with Ly6G$^+$ cells (red), CD31$^+$ cells (green) and SHG (second harmonic generation) (blue). Right images show spot and surface transformed data with spot (Ly6G$^+$) and surface (CD31$^+$). **(G)** Representative images of the transformed 3D-intravital microscopy of neutrophil (left) and random spot (right) proximity distribution to blood vessel in the 2d IMQ-skin, scale bar=50μm. **(H)** Bar graphs showing the comparative distribution of neutrophils and random spots to the blood vessel in the IMQ-skin at 0d (upper) and 1d of IMQ treatment (lower) (n=3 for each). Data are pooled from ≥2 independent experiments with each dot in the graphs representing one experimental subject. For 3D-intravital microscopy, experiment was done in 1 subject per group with each dot in the graphs representing individual acquired tissue plane. Data are shown as mean ± S.E. *$p$<0.05; **$p$<0.01; ***$p$<0.001; ****$p$<0.0001.

**Figure S2 Overactive neutrophils have pathological function in psoriasis.**

**(A)** Gene ontology analysis of neutrophils from Vas-treated (n=3) and IMQ-reated skin (n=3). **(B)** Representative FACS plot and quantification of peripheral blood neutrophils at 4d from IgG or anti-Ly6G antibody injected IMQ-induced mice (n=3 for each). **(C)** Clinical examination of dorsal IMQ-skin at 0-4d with IgG or anti-Ly6G treatment. Upper: representative skin images, lower: erythema and scaling scoring (IgG: n=4, anti-Ly6G: n=4). Data are pooled from ≥2 independent experiments with each dot shown in the bar graph represents data from each individual and shown as mean ± S.E. For RNA-seq, analysis was done to the data pooled from ≥2 replicates. *$p$<0.05.

**Figure S3 Psoriasis induces expansion of L86K-defined HSPC in BM.**

**(A)** FACS plots of LSK (Lin$^-$ Sca-1$^+$c-Kit$^+$) (upper) and L86K (Lin$^-$CD86$^+$c-Kit$^+$) cells (lower) in BM at 16h, 2d, and 4d after Vas or IMQ treatment (n=3-6 for each). **(B)** FACS plot of immunophenotypic lt-HSC, st-HSC, MPP2, MPP3, and MPP4 defined by L86K (S3A, lower panel) in the bone marrow of mice treated with Vas/IMQ (n=3-6 each). **(C)** Graphs depicting the absolute number of LSK (left), L86K (middle) and L86K-defined HSPC (lt-HSC, st-HSC, MPP2-4) (right) in the bone marrow of mice treated with Vas/IMQ (n=3-6 for each). **(D)** Pie charts depicting relative composition of hematopoietic cells ($CD45^+$) in BM (upper) and skin (lower) at 16h, 2d, and 4d after Vas or IMQ treatment (n=3-6 each). Data are pooled from ≥2 independent experiments. Each dot shown in the bar graph represents data from each individual and shown as mean±S.E. **p*<0.05; ***p*<0.01; ****p*<0.001; *****p*<0.0001.

**Figure S4 IL-1 signals do not contribute to psoriasis-driven emergency granulopoiesis.** **(A)** Experimental scheme for (B): Vaseline (Vas) or Imiquimod (IMQ) topically applied to tohe dorsal skin daily for 4d and serum analysis was conducted at 4d. **(B)** Cytokines protein measurement from the sera (Vas: n=4, IMQ: n=5). **(C)** Experimental scheme of IMQ-induced psoriasis (0-3d) with daily PBS/Anakinra intrapertoneal injection (from -1d to 3d) and analyzed at 4d for results depicted in (D-H). **(D)** Representative FACS plots of immature ($Ly6G^{lo}$) and mature ($Ly6G^{hi}$) granulocytes pregated on $CD45^+$ BM cells (left). The proportion of $Ly6G^{lo}$ and $Ly6G^{hi}$ cells (n=4 for each) (right). **(E)** Representative FACS plots of myeloid cell fractions in the dorsal skin preagated on $CD45^+CD11b^+$ cells (left). Cell number quantification graph of the IMQ-induced mice treated with PBS/Anakinra (n=4 for each) (right). **(F)** Skin clinical examination upon psoriasis disease course. Representative naked skin images with scale bar=1cm (left upper panel) and skin histology by HE staining with black line indicates 100μm (left lower panel) Daily clinical scoring (middle) and epidermal thickness measurment of IMQ-induced psoriatic skin following treatment with PBS/Anakinra (n=3-4 for each). Data are pooled from ≥2 independent experiments with each dot shown in the bar chart represents individual subject and shown as mean±S.E. **p*<0.05.

**Figure S5 G-CSF neutralization reduces circulating neutrophils, hematopoietic activation in bone marrow, and psoriasis clinical manifestation.** **(A)** Representative FACS plot along with the proportional and absolute cell number of circulating phenotypic neutrophils bar chart in IMQ-induced psoriasis mice treated with IgG/anti-GCSF (n=3/group). **(B)** FACS analysis of the $Lin^-c\text{-}Kit^+Sca\text{-}1^+$ (LSK) in the bone marrow of IMQ-induced psoriasis treated with IgG/anti-G-CSF (n=3 each). **(C)** Psoriasis clinical manifestation of the IMQ-induced mice which was monitored daily at the naked dorsal skin (upper) and clinically (erythema and scaling) graphed (lower) with i.p. injection of IgG/anti-G-CSF (n=3 for each). Data are pooled from ≥2 independent experiments with each dot shown in the chart represents the measurement from 1 experimental subject and shown as mean±S.E. **p*<0.05; ***p*<0.01; ****p*<0.001; *****p*<0.0001.

**Figure S6 Clustering and relative transcript measurement of human psoriasis skin biopsies. (A)** UMAP clustering of RNA-seq data from healthy and psoriatic human skin biopsy (normal: n=82, psoriasis: n=92). **(B)** Relative expression of dermatitis-related cytokines (normal: n=82, psoriasis: n=92). **(C-F)** Reanalysis of single cell RNA-seq dataset GSE173706 (normal: n=8, psoriasis: n=14) consisting of UMAP clustering results **(C),** gene list for clusters annotation **(D)**, cell frequency counting on each annotated cell types **(E)**, and re-clustering of annotated endothelial cell subset **(F)**. Each dot shown in the chart represents the measurement from 1 experimental subject and shown as mean±S.E. $^{*}p<0.05$; $^{**}p<0.01$; $^{***}p<0.001$; $^{****}p<0.0001$.

**SUPPLEMENTAL MATERIAL**

**Table for reagent or resource**

| REAGENT or RESOURCE | SOURCE | IDENTIFIER |
|---|---|---|
| **FACS antibodies** | | |
| Biotin anti-CD3e | Biolegend | 100304 |
| Biotin anti-CD8a | Biolegend | 100704 |
| Biotin anti-CD4 | Biolegend | 100404 |
| Biotin anti-NK1.1 | Biolegend | 108704 |
| Biotin anti-IL7ra | Biolegend | 121104 |
| Biotin anti-Ter119 | Biolegend | 116204 |
| Biotin anti-Gr1 | Biolegend | 108404 |
| Biotin anti-CD11b | Biolegend | 101204 |
| Biotin anti-B220 | Biolegend | 103204 |
| anti-CD16/32 | Biolegend | 101302 |
| Brilliant Violet 605 Streptavidin | Biolegend | 405229 |
| Brilliant Ultra Violet 395 CD45 | BD | 564279 |
| BV421 CD45 | Biolegend | 103133 |
| FITC CD45.2 | Biolegend | 109806 |
| FITC CD34 | invitrogen | 11-0341-85 |
| Pacific Blue CD48 | Biolegend | 103418 |
| APC ckit (CD117) | Biolegend | 105812 |
| PE CD150 | Biolegend | 115904 |
| PE-Cy7 CD11c | Biolegend | 117318 |
| PE-Cy7 CD101 | eBioscience | 25-1011-80 |
| BV605 MHC2 | Biolegend | 107639 |
| BV421 CD31 | Biolegend | 102423 |
| APC CD140a | Biolegend | 135908 |
| PE CD49f | Biolegend | 313611 |
| PE-Cy5 CD135 | Biolegend | 135312 |
| APC-Cy7 Sca-1 | Biolegend | 108126 |
| Brilliant Violet 785 Ly6G | Biolegend | 127645 |
| PE-Cy7 Ly6G | Biolegend | 127618 |
| Brilliant Violet 711 Ly6C | Biolegend | 128037 |
| APC-Cy7 CD11b | Biolegend | 101226 |

| | | |
|---|---|---|
| PE Langerin (CD207) | Biolegend | 144204 |
| InVivoMAb rat IgG2 isotype control, anti-trinitrophenol | Bio X Cell | BE0089 |
| InVivoMAb anti-mouse Ly6G | Bio X Cell | BE0075-1 |
| Mouse IgG antibody | R&D systems | AF007 |
| Mouse G-CSF antibody | R&D systems | MAB414 |
| **Chemicals, peptides, and recombinant proteins** | | |
| Hair removing cream | Veet | PM8220626 |
| Vaseline cream (100% White Petrolatum) | Unilever | 521234500 |
| Beselna Cream 5% (Imiquimod) | Mochida Pharmaceutical Co., Ltd. | MO651 |
| KINERET (Anakinra) | Biovitrum | 49163-54388 |
| AmbionTM Nuclease-Free Water | Invitrogen | AM9937 |
| Hanks' Balanced Salt Solution 10x | Sigma | H4641-500ML |
| RPMI-1640 with L-Glutamine and Phenol Red | Wako | 189-02025 |
| Ethanol 99.5% | Fujifilm | 052-03343 |
| LiberaseTM | Roche | 5401127001 |
| DNAse1 | Roche | 10104159001 |
| DNAse I Amplification grade | Invitrogen | 10977015 |
| 1 mol/L-HEPES Buffer solution | Nacalai Tesque | 17557-94 |
| Mayer's Hematoxylin solution | Wako | 131-09665 |
| Eosin Y (Sodium Tetrabromofluorescein) | Fujifilm | 058-00062 |
| Xylene | Fujifilm | 241-00091 |
| Paraformaldehyde | Fujifilm | 1613-20145 |
| Entellan TM | Sigma | 1.07961.0100 |
| Bovine Serum Albumin | Merck | A9418-5G |
| RNEasy Mini Kit | Qiagen | 74104 |
| THUNDERBIRD SYBR qPCR/RT set | Toyobo | QPS201 |
| Primescript RT Master Mix | Takara Bio | RR036A |
| Hoechst 33342, trihydrochloride | Life Technologies | H3570 |
| Propidium Iodide | Sigma | P4170-10MG |
| LPS-EB Ultrapure | Invivogen | tlrl-3pelps |
| Imiquimod | Sigma | I5159-200MG |
| Gelatin from cold fish skin | Merck | G7041-100G |
| DMEM High Glucose | Wako | 044-29765 |

| | | |
|---|---|---|
| CultureSure DMSO | Wako | 031-24051 |
| Non-Essential Amino Acids Solution | Gibco | 11140050 |
| GlutaMAX™ Supplement | Gibco | 35050-061 |
| Penicillin-Streptomycin | Wako | 164-25251 |
| Fetal Bovine Serum | Nichirei | 174012-500ML |
| Mouse G-CSF ELISA Kit | Proteintech | KE10025 |
| LEGENDplex™ Inflammation Panel (13-plex) with Filter Plate | Biolegend | 740150 |
| NP40 | Thermo Fischer | 28324 |
| RNAsein plus | Promega | N2611 |
| Real Time Cell Lysis buffer | Roche | 06366821001 |
| 5x RT Buffer | Takara | RR037A |
| ERCC RNA Spike-In Mix | Thermo | 4456740 |
| 20x enzyme mix | Takara | RR037A |
| T4 gene 32 protein | NEB | M0300L |
| Oligo(dT) 18 primer | Thermo | SO131 |
| $1^{st}$ NSR primer mouse | SIGMA | N/A |
| $2^{nd}$ NSR primer mouse | SIGMA | N/A |
| NEB buffer 2 | NEB | M0212L |
| Klenow fragment (3'-5' exo-) | NEB | M0212L |
| dNTPs | Takara | 4030 |
| AMPure XP beads | Beckman Coulter | A63881 |
| Tagment DNA buffer | Illumina | FC-131-1096 |
| Index primers | Illumina | FC-131-2001 |
| High Sensitivity D5000 Screen Tape | Agilent | 5067-5592 |
| GenNext® NGS Library Quantification kit | Toyobo | NLQ-101 |
| **Experimental models: Organisms/strains** | | |
| *Mus musculus* (C57BL/6JmsSlc) | Japan SLC,Inc | N/A |
| **Software and algorithms** | | |
| FACSDiva Software | BD | N/A |
| Flowjo 10.10.0 | BD | N/A |
| GraphPad Prism 10.1.2 (324) | Dotmatics | N/A |
| ImageJ 1.53t | NIH | N/A |
| R Studio build 394 | R | N/A |

| | | |
|---|---|---|
| **RNA-seq data** | | |
| Human RNA-seq data (Journal of Investigative Dermatology) | PMID: 24441097 | https://www.ncbi.nlm.nih.gov/geo/query/acc.cgi?acc=GSE54456 |
| Human RNA-seq data (Nature Communications) | PMID: 37308489 | https://www.ncbi.nlm.nih.gov/geo/query/acc.cgi?acc=GSE173706 |
| RNA-seq data of ouse skin neutrophils | PRJDB37718 | https://ddbj.nig.ac.jp/search |

N/A, not applicable

**Primer list for RT-PCR**

| Gene(s) | 5' -> 3' | RT-PCR Oligonucleotides |
|---|---|---|
| *Il17a* | Foward | CTGCTGAGCCTGGCGGCTAC |
| | Reverse | CATTGCGGTGGAGAGTCCAGGG |
| *Il17f* | Foward | ACCCGTGAAACAGCCATGGTCAAG |
| | Reverse | CCCATGGGGAACTGGAGCGG |
| *Il23* | Foward | TCCTCCAGCCAGAGGATCACCC |
| | Reverse | AGAGTTGCTGCTCCGTGGGC |
| *Il22* | Foward | CAGCTCCTGTCACATCAGCGGT |
| | Reverse | AGGTCCAGTTCCCCAATCGCCT |
| *Tnfa* | Foward | GCCCACGTCGTAGCAAACCAC |
| | Reverse | GCAGGGGCTCTTGACGGCAG |
| *Il6* | Foward | CCTCTCTGCAAGAGACTTCCAT |
| | Reverse | AGTCTCCTCTCCGGACTTGT |
| *Il1a* | Foward | CTGGAAGAGACCATCCAACCC |
| | Reverse | ACTTCCTGTTGCAGGTCATTT |
| *Csf3* | Foward | GCTTCCTGCTTAAGTCCCTG |
| | Reverse | CTTAGGCACTGTGTCTGCTG |
| *Cxcl1* | Foward | CCGAAGTCATAGCCACACTCAA |
| | Reverse | GCAGTCTGTCTTCTTTCTCCGTTA |
| *Cxcl2* | Foward | GAAGTCATAGCCACTCTCAAGG |
| | Reverse | CCTCCTTTCCAGGTCAGTTAGC |

| | | |
|---|---|---|
| *Cxcl5* | Foward | TGATCGCTAATTTGGAG GTGAT |
| | Reverse | TAGCTTTCTTTTTGTCACTGCC |
| *Ccl2* | Foward | TCTGGGCCTGCTGTTCACA |
| | Reverse | GGATCATCTTGCTGGTGAATGA |
| *Tlr7* | Foward | GTTCTTGACCTTGGCACTA |
| | Reverse | CCGTGCATATTCATCGTA |
